# Metabolic cross-feeding allows a gut microbial community to overcome detrimental diets and alter host behaviour

**DOI:** 10.1101/821892

**Authors:** Sílvia F. Henriques, Lúcia Serra, Ana Patrícia Francisco, Zita Carvalho-Santos, Célia Baltazar, Ana Paula Elias, Margarida Anjos, Tong Zhang, Oliver D. K. Maddocks, Carlos Ribeiro

## Abstract

The impact of commensal bacteria on the host arises from complex microbial-diet-host interactions. Mapping metabolic interactions in gut microbial communities is therefore key to understand how the microbiome influences the host. Here we use an interdisciplinary approach including isotope-resolved metabolomics to show that in *Drosophila melanogaster, Acetobacter pomorum* (*Ap*) and *Lactobacillus plantarum* (*Lp*) establish a syntrophic relationship to overcome detrimental host diets and identify *Ap* as the bacterium altering the host’s feeding decisions. Specifically, we show that *Lp* generates lactate which is used by *Ap* to produce and provide amino acids that are essential to *Lp* allowing it to grow in imbalanced diets. Lactate is also necessary and sufficient for *Ap* to alter the fly’s protein appetite. Our data show that gut bacterial communities use metabolic interactions to become resilient to detrimental host diets and to ensure the constant flow of metabolites used by effector bacteria to alter host behaviour.

## Introduction

In all organisms, including humans, diet is a critical determinant of health and wellbeing (Simpson and Raubenheimer, 2012). As the building blocks of proteins, dietary amino acids (AAs) play a pivotal role in determining the fitness of all animals. Because they cannot be efficiently synthesized, essential amino acids (eAAs) need to be acquired through the diet (Mccoy *et al.*, 1935; Rose, 1957). Furthermore, universally, over-ingestion of dietary AAs shortens lifespan and negatively impacts healthspan (Grandison *et al.*, 2009; Levine *et al.*, 2014; Solon-Biet *et al.*, 2014). Given the importance of a balanced dietary intake of AAs, organisms are able to direct their feeding choices to homeostatically compensate both for the lack and over-ingestion of AAs (Gosby *et al.*, 2011; Leitão-Gonçalves *et al.*, 2017; Ribeiro and Dickson, 2010; Simpson *et al.*, 2015; Solon-Biet *et al.*, 2019). In *Drosophila melanogaster* females, both AA deprivation and mating induce changes in specific neuronal circuits which modulate food choice, leading to a drastic increase in protein appetite, (Ribeiro and Dickson, 2010; Steck *et al.*, 2018). Remarkably, the removal of any of the ten eAAs from the fly diet is sufficient to induce this strong protein appetite (Leitão-Gonçalves *et al.*, 2017). While much is known about the physiological and neuronal processes underlying bulk food intake, less is known about how nutrient specific appetites are controlled. Given the importance of a balanced diet and especially of a balanced intake of dietary AAs for animal fitness it is key that we advance our understanding about the factors controlling nutritional choices.

The gut microbiome has emerged as an important modulator of host physiology and behaviour (Clemente *et al.*, 2012; Cryan *et al.*, 2019; Martino *et al.*, 2017; Subramanian *et al.*, 2015; Vuong *et al.*, 2017). As such, gut bacteria have also been shown to influence feeding behaviour and food choice (Breton *et al.*, 2016; Leitão-Gonçalves *et al.*, 2017; Vijay-Kumar *et al.*, 2010). A key challenge of current microbiome research is to identify the mechanisms underlying the effect of the microbiome on the host. While in specific cases single microbes can be identified as the sole drivers affecting the host (Oh *et al.*, 2019; Schretter *et al.*, 2018; Shin *et al.*, 2011; Storelli *et al.*, 2011; Yoon *et al.*, 2016), in the majority of cases it is clear that gut microbes act on the host as a community rather than as isolated biotic factors (Gould *et al.*, 2018; Olson *et al.*, 2018; Rekdal *et al.*, 2019). This is likely also the case for the impact of the gut microbiome on the brain. In mice for example, *Akkermansia* and *Parabacteroides* bacteria have been suggested to act together to mediate the beneficial effects of the ketogenic diet in seizure prevention (Olson *et al.*, 2018). Conceptually, two main mechanisms can explain why only a community of microorganisms can affect the physiology and behaviour of the host: 1) the need for the tandem catalytic activity of enzymes from different microbiota for the production of metabolites acting on the host (Rakoff-Nahoum *et al.*, 2016; Sonnenburg *et al.*, 2010) and/or 2) the need for an obligatory mutualistic metabolic relation (syntrophy) to sustain the growth of specific microorganisms so that they can promote the observed physiologic effect in the host (Morris *et al.*, 2013). To identify the mechanisms by which bacterial communities act on the host we will need to map out the relevant interactions among gut microbes influencing the host and identify the molecular and metabolic mechanisms by which they do so. This remains a daunting task given the large number of microbial species constituting vertebrate gut microbiomes.

Host diet is considered one of the most relevant determinants of human gut microbiome variation (David *et al.*, 2014; Johnson *et al.*, 2019; Koenig *et al.*, 2011). The microbiome composition changes rapidly in response to new food choices, such as shifting from plant-based to animal-based diets (David *et al.*, 2014; Dhakan *et al.*, 2019; Johnson *et al.*, 2019) or changes in the protein to carbohydrate intake ratio (Holmes *et al.*, 2017). Adding to this complexity, in humans the impact of diet on the microbiome is highly personalized (Johnson *et al.*, 2019). While in vitro experiments have started to systematically disentangle the nutritional preferences of single human gut microbes (Tramontano *et al.*, 2018), we are very far from achieving a coherent mechanistic picture of how bacterial dietary needs shape the microbiome and its capacity to influence the host. Given the large number of nutrients required by the host and the nutritional complexity of natural foods, identifying how single nutrients affect the microbiome and hence the host, remains a key challenge in current microbiome research.

Since the microbiome serves as a stable regulatory factor contributing to host physiology, it has been proposed that it should be resilient to dietary perturbations (Greenhalgh *et al.*, 2016; Lozupone *et al.*, 2012). As any other organism, gut microbes have very different nutritional requirements which at the individual species level, are largely defined by the biosynthetic capacities encoded at the genome level. The complex metabolic interactions within microbial communities can however profoundly alter the impact of nutrients on the physiology of the different species in the community (Ponomarova and Patil, 2015). In the context of the gut microbiome this could contribute to the emergence of dietarily resilient communities which would be stable even if the host diet lacks nutrients which are essential for specific members of the community. Identifying the mechanisms allowing gut microbe communities to overcome dietarily challenging conditions could therefore significantly expand our understanding of the conditions in which the gut microbiome becomes susceptible to changes in host diet. This knowledge could also be used to guide the development of tailored interventions aiming at strengthening the resilience of gut microbe communities, thereby ensuring their continuous beneficial impact.

*Drosophila melanogaster* has emerged as a powerful, experimentally tractable system to identify the mechanisms by which gut microbes interact with the host to influence diverse traits ranging from metabolism, to growth and behaviour (Douglas, 2018; Martino *et al.*, 2017). The adult *Drosophila* has a simple microbiome, typically containing less than 20 taxa, and mainly populated by species from the *Acetobacter* and *Lactobacilli* genera which can be cultivated in the laboratory and are amenable to genetic manipulations (Broderick and Lemaitre, 2012; Wong *et al.*, 2011). It is furthermore easy to generate and maintain germ-free animals as well as to reconstitute gnotobiotic animals with a predefined microbiome. Importantly, its powerful genetic and genomic toolset, the ability to perform large-scale, hypothesis-agnostic screens, and the availability of a chemically defined diet (Piper *et al.*, 2017, 2014) make the fly an ideal system to identify core mechanistic principles governing diet-host-microbiome interactions (Baenas and Wagner, 2019; Douglas, 2018).

In this study we show that a syntrophic relation between *Ap* and *Lp*, two abundant strains making up the fly microbiome, is at the base of their ability to suppress yeast appetite in flies deprived of eAAs. Using a chemically defined fly diet we found that *Ap* is able to promote *Lp* growth in flies reared in media lacking isoleucine (Ile), an eAA for *Lp*. To explore the impact of host dietary conditions on the bacterial community we adapt the fly holidic medium to be able to grow bacteria in vitro in high throughput. By combining this diet with isotope resolved metabolomics, we show that the presence of *Lp* stimulates *Ap* to produce and excrete Ile as well as other AAs into the media. These can then be used by *Lp* to grow in the absence of those AAs in the diet. We furthermore identify lactate as the main contribution of *Lp* to the bacterial mutualistic relationship. As such it is possible to substitute *Lp* by lactate and observe the same level of protein appetite suppression, showing that this metabolite is necessary and sufficient for modifying protein appetite in the presence of *Ap*. Interestingly, lactate was required for *Ap* to produce Ile, and using stable isotope labelled lactate we show that it serves as a major precursor for the synthesis of AAs by *Ap*. These data provide clear evidence of a “circular economy” in which *Lp* derived lactate is used by *Ap* to generate AAs which allow *Lp* to proliferate and provide lactate to the community. Given that *Ap* is sufficient to modify the behaviour of the host and is able to synthesize all eAAs, we tested the hypothesis that *Ap* modifies host behaviour by replenishing AAs in the malnourished host. We, however provide multiple evidence contradicting this hypothesis. We found that the bacteria only secrete extremely low levels of Ile compared to the levels required to alter behaviour in physiological conditions. We also show that while the *Ap*/*Lp* community is beneficial for the host, allowing it to increases egg laying in malnourished females, *Ap* combined with lactate does not increase egg laying in these females. This allows us to functionally separate the beneficial effect of the community on egg laying from the effect on behaviour, and strongly suggests that some other *Ap*-derived factor than bacterial produced eAAs influences food choice. This work demonstrates the importance of bacterial communities as the relevant explanatory unit necessary to understand how the gut microbiome impacts the host. We furthermore uncover the molecular basis of a circular metabolic cross-feeding relationship that supports the stability of a gut microbial community making it resilient to the nutritional environment that the host may encounter. The resilience to dietary challenges allows the community to exert its beneficial impact when the animal is malnourished and ensures the continuous availability of metabolic precursors used by the behavioural effector species to alter brain function.

## Results

### A gut bacterial community consisting of *A. pomorum* and *L. plantarum* buffers yeast appetite in flies deprived of essential amino acids

The impact of the microbiome on host physiology mostly emerges from the complex interaction of diet, gut bacteria and host physiology. The use of chemically defined (holidic) diets in combination with gnotobiotic animals are ideal tools to disentangle these complex interactions (Leitão-Gonçalves *et al.*, 2017; Piper *et al.*, 2014). We pioneered the use of this approach to show that in mated *Drosophila melanogaster* females, the removal of any of the ten eAA, induces a strong and specific appetite for yeast (the main protein source for flies) (Figure 1A and Figure S1A) (Leitão-Gonçalves *et al.*, 2017). Here and throughout this study we decided to focus on two eAAs, Ile and histidine (His), as proxies for all ten eAAs. These two AAs represent very different chemical and biological classes of eAAs with one being a branched chain amino acid (BCAA), allowing us to cover a broad spectrum of biological activities. We used this framework to explore the interaction of dietary eAAs and the gut microbiome on protein appetite. Remarkably, a community of two bacteria, consisting of *Lactobacillus plantarum* (*Lp*) together with *Acetobacter pomorum* (*Ap*), can significantly suppress the yeast appetite of flies deprived of eAAs (Figure 1A and Figure S1A). Interestingly, the presence of any of these bacteria alone cannot suppress protein appetite (Leitão-Gonçalves *et al.*, 2017). These results clearly show that the impact of the microbiome on feeding decisions relies on the presence of a minimal bacterial community consisting of only two members. This is in line with the common observation that many effects of the microbiome on host physiology and behaviour can only be understood at the level of microbial communities and not single bacteria (Gould *et al.*, 2018; Leitão-Gonçalves *et al.*, 2017; Olson *et al.*, 2018; Rekdal *et al.*, 2019).

**Fig. 1.**
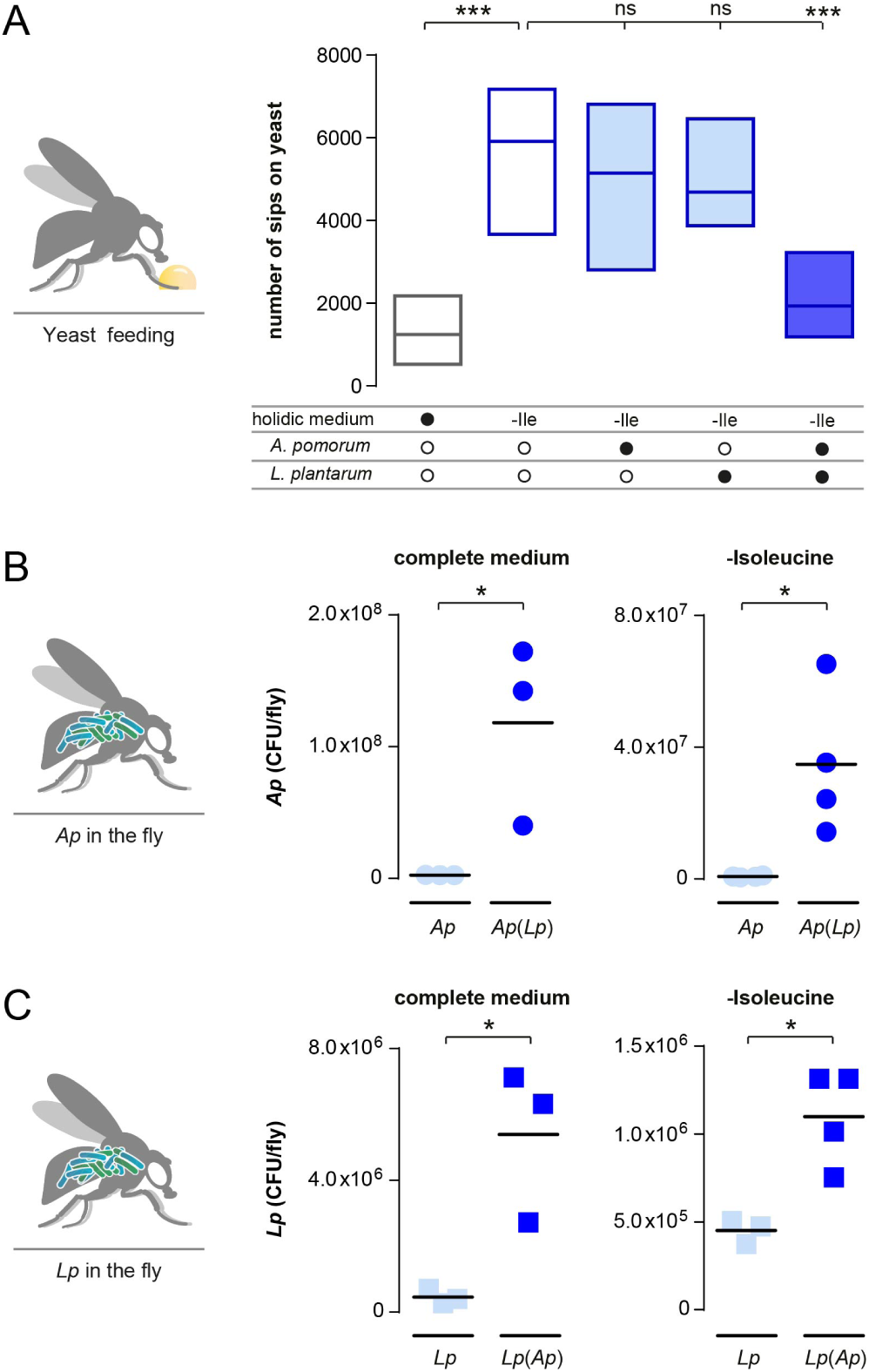
*Ap* and *Lp* biassociation reduces yeast appetite and promotes gut bacterial growth independent of dietary Isoleucine content. (A) Number of sips on yeast of flies kept in complete holidic medium or medium without Ile (-Ile) that were monoassociated (light blue) with *Ap* or *Lp*, or biassociated (blue) with both commensal species. In this and other Figures, empty boxes represent germ-free conditions and filled boxes represent bacteria-associated conditions. Boxes represent median with upper and lower quartiles. n = 25-35. Significance was tested using a Kruskal–Wallis test with a Dunn’s multiple comparison test. Filled black circles represent a complete holidic medium or association with specific bacteria. Open black circles represent the absence of specific bacteria. (B and C) Number of *Ap* (circles, B) and *Lp* (squares, C) colony forming units (CFUs) from extracts of monoassociated (light blue) and biassociated (dark blue) flies kept in complete holidic media or medium lacking Ile (-Isoleucine). *Ap* and *Lp* labels data points representing the number of CFUs detected in monoassociated flies and *Ap*(*Lp*) and *Lp*(*Ap*) the measurement of *Ap* or *Lp* in biassociated flies. The black line represents the mean. n=3-4. Significance was tested using an unpaired t-test. Not significant (ns) p > 0.05, * p 0.05, *** p 0.001.

### The bacterial community buffers single bacteria from adverse effects of imbalanced host diets

To mechanistically understand why the behavioural impact of the gut microbiome requires a community, we decided to start by testing the impact of diet and community composition on the titers of the different bacteria in the gut of the host. Diet is a potent modulator of the microbiome, and it is very possible that the composition of the different fly diets may impact bacterial composition and hence their effect on the host (De Filippo *et al.*, 2010; Turnbaugh *et al.*, 2009; Wang *et al.*, 2014; Wong *et al.*, 2014). Especially as both Ap and Lp are auxotrophic for different nutrients including eAAs (Saguir and de Nadra, 2007; Teusink *et al.*, 2005). We therefore compared the internal bacterial load of flies inoculated with single cultures or co-cultures of *Ap* and *Lp* maintained for three days on a complete holidic medium, or in media lacking Ile or His. In the tested flies the number of both *Ap* and *Lp* cells was consistently higher in the co-culture conditions when compared to the bacterial load in flies inoculated with each of the bacteria alone (Figure 1B and 1C and Figure S1B and S1C). This is in agreement with other reports showing synergistic effects in bacterial growth of Acetobacter and Lactobacillus species in *Drosophila* (Newell and Douglas, 2014; Sommer and Newell, 2019).

Interestingly, the increase in bacterial load when the microbes were co-cultured was observed in the complete medium and the media lacking Ile or His (Figure 1B and 1C and Figure S1B and S1C). This is expected for the condition in which we removed His from the diet, as both *Ap* and *Lp* should be able to synthesize this AA (Martino *et al.*, 2016; Newell *et al.*, 2014; Saguir and de Nadra, 2007; Shin *et al.*, 2011). However, given that *Lp* is not supposed to synthesize Ile it is unexpected that in flies maintained on media lacking Ile, the levels of *Lp* can increase (Figure 1C). This clearly indicates that in the co-culture condition *Lp* can grow in media lacking Ile, which given the predictions from the *Lp* genome and previous in *vitro* growth experiments should not be possible (Martino *et al.*, 2016; Newell *et al.*, 2014; Saguir and de Nadra, 2007). These data indicate that the presence of *Ap* allows *Lp* to overcome its Ile auxotrophy and that the community is therefore able to buffer single bacteria from adverse effects of host diets lacking specific essential nutrients.

### *Ap* allows *Lp* to overcome its isoleucine auxotrophy

In order to carefully dissect the impact of the diet on bacterial growth and exclude the host as a confounding factor, we adapted the holidic fly medium to grow bacteria in *in vitro* liquid cultures (see material and methods). This also allowed us to perform bacterial dietary manipulations in large scale while maintaining the bacteria in the diet of the host. We first cultivated *Ap* and *Lp* alone in the liquid versions of the complete holidic fly medium or media lacking either Ile or His and assessed the growth of these bacteria over three days. *Ap* grew to the same extent in complete medium, and in media lacking His or Ile (Figure 2A and Figure S2A). This confirms the data obtained in the host and the known biosynthetic activities of this bacterium (Figure 1B and Figure S1B). It also shows that the liquid version of our holidic medium is suitable for cultivating *Drosophila* commensal bacteria. Interestingly, in the *in vitro* situation we did not observe a clear increase in growth in the co-culture condition when compared to the condition in which we grew *Ap* alone (Figure 2A and Figure S2A vs. Figure 1B and 1C). This strongly suggests that the overall increase in commensal bacteria proliferation observed in the host when they grow as a community, is specific to the intestinal niche. This points to beneficial effects of the community which are specific to the situation in which bacteria are growing in the animal gut and highlights the strength of the *in vitro* culture system in isolating pure dietary effects from effects resulting from bacteria-host interactions.

**Fig. 2.**
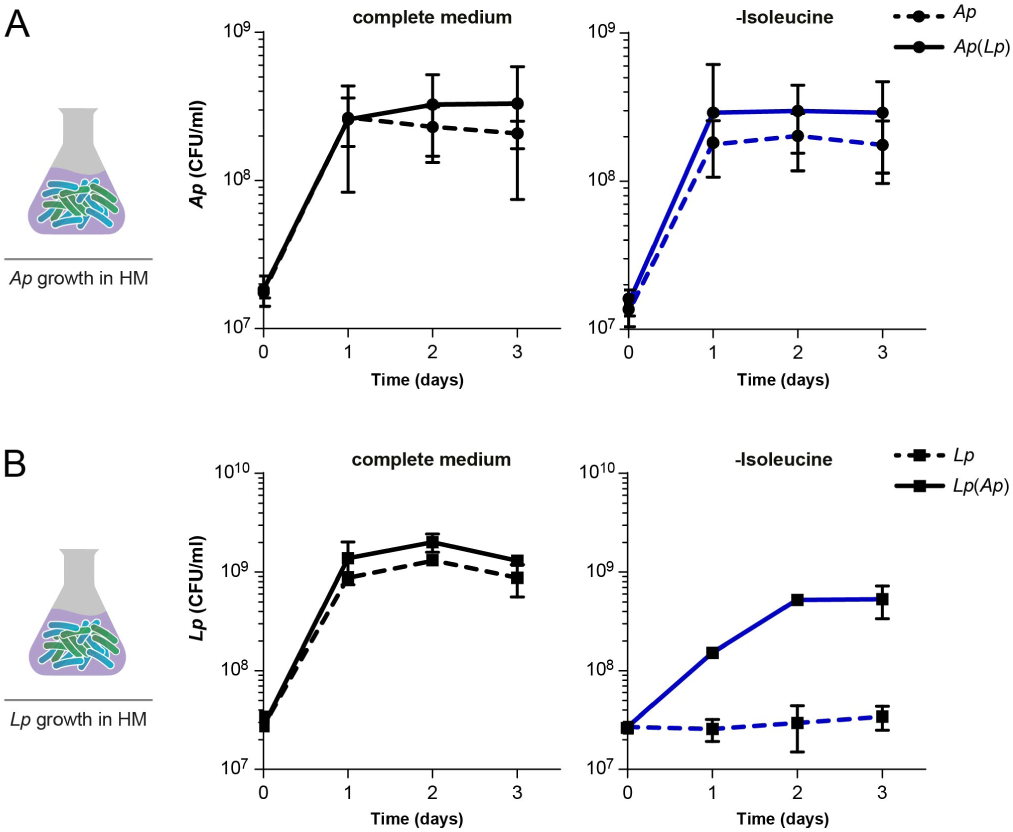
*In vitro* growth measurements show that *Lp* can overcome its auxotrophy for Isoleucine when in co-culture with *Ap*. In vitro growth curves of *Ap* (A) and *Lp* (B) in complete liquid holidic medium (black line) or in liquid medium with-out Ile (blue line) plotted as CFUs at different time intervals. *Ap* and *Lp* (dashed lines) are data points representing the number of CFUs detected in monoassociated flies and *Ap*(*Lp*) and *Lp*(*Ap*) (continuous lines) the measurement of *Ap* or *Lp* in co-culture conditions. The number of CFUs per ml of culture was determined by collecting and cultivating liquid culture samples in selective media at the indicated time points (as described in material and methods). Data points represent mean values of three to four biological replicates and error bars the standard deviation of the mean.

While there was no effect of removing His from the medium on the growth of *Lp* alone (Figure S2B), *Lp* could not grow in the absence of Ile (Figure 2B). However, when co-cultured with *Ap, Lp* was able to efficiently grow in media lacking this AA despite being auxotrophic for it (Figure 2B). The ability of *Ap* to support *Lp* growth in liquid medium lacking Ile, reproduces the observations made in the host situation. Our findings strongly suggest that *Ap* does so by providing Ile to *Lp*.

### Isoleucine and other amino acids are synthesized by the bacterial community

Our *in vivo* and *in vitro* bacterial growth data suggest that *Ap* would have to synthesize and secrete Ile to allow *Lp* to overcome the dietary lack of this eAA. To test this hypothesis, we decided to use stable isotope labelling to measure *de novo* synthesis and secretion of AAs from dietary glucose by the gut bacteria. First we cultivated *Ap* alone in -Ile media containing uniformly labelled ^13^C_6_-D-glucose to track the synthesis of _13_C-Ile and of other _13_C-labeled AAs. Given our interest in the secreted fraction of AAs we measured the amount of labelled AAs in the supernatant of the cultures using liquid chromatography mass spectrometry (LC-MS). While *Ap* should be synthesizing Ile to be able to grow in a medium lacking this eAA, we could not detect an increase in secreted labelled Ile when we cultivated *Ap* alone (compared to both the ^12^C_6_-D-glucose no-labelling control and the no-growth control (0h)) (Figure 3). However, when *Ap* grew in a community with *Lp*, we could detect an increase in the presence of multiple ^13^C-labelled isotopomers (m+3, m+4, m+5) of Ile when compared to the controls and the *Ap* single culture (Figure 3). We did not detect an increase in secreted, synthesized Ile when the bacteria were grown in media that contained Ile (either complete medium or medium lacking His), suggesting that the absence of dietary AAs stimulates the biosynthesis of this metabolite (Figure S3). Furthermore, when analysing other AAs we could see an overall increase in labelled and secreted AAs in the co-culture condition when compared to when *Ap* was grown in isolation. (Figure S4). Importantly, the same pattern of AA labelling was observed in 48h cultures with the expected exception that heavier isotopomers were also detected, attesting to the robustness of our findings (Figure S5). These results strongly suggest that the production of AAs by *Ap* and/or their accumulation in the medium is stimulated by the presence of *Lp*. In conclusion, these data show that *Ap* produces and secretes Ile and other AAs when growing in co-culture with *Lp*, providing the biochemical basis by which the bacterial community overcomes nutritional challenges posed by imbalanced host diets.

**Fig. 3.**
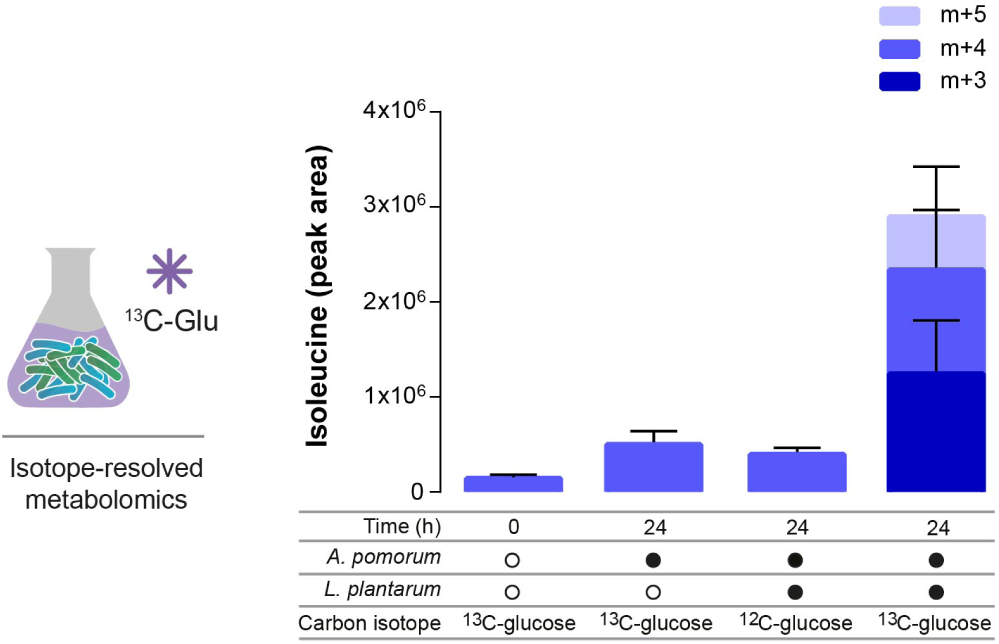
*Ap* synthesizes and secretes Isoleucine in the presence of *Lp*. The stacked bars represent the relative amount of Ile isotopomers measured in the supernatant of liquid holidic medium of bacterial cultures lacking Ile and either containing unlabelled glucose (^12^C-glucose control), or uniformly labelled ^13^C-glucose (^13^C-glucose). Heavy labelled Ile isotopomers were measured using LC-MS in samples collected after 24h of bacterial growth and displayed as metabolite peak area. No bacteria were added to the media representing time “0”. The number of heavy carbons incorporated per Ile molecule is indicated as m+n, where n = the number of ^13^C. Filled black circles represent the presence of specific bacteria in the culture. Open black circles represent the absence of specific bacteria. Each data point represents the mean of three biological replicates and the error bars represent the standard error of the mean for each isotopomer.

### *Lp* contributes to the bacterial syntrophy through the production of lactate

We have shown that *Ap* allows *Lp* to overcome its Ile auxotrophy by providing this limiting AA. Furthermore, our data clearly show that both the behavioural effect on the host as well as the increase in available AAs is a community effect depending on the presence of *Lp*. Therefore, *Lp* must be making a critical contribution to the community both in the host as well as when growing *in vitro*. What could be the form of this contribution? We have shown that both *Lp* and *Lactobacillus brevis* are interchangeable in their capacity to suppress yeast appetite when inoculated in co-culture with *Ap* (Leitão-Gonçalves *et al.*, 2017). Therefore, given that both are lactic acid-producing bacteria, one attractive hypothesis is that *Lp* provides lactate to *Ap* which then is used as a metabolic precursor by this bacterium. Supporting this hypothesis in the labelled metabolomics data we could detect significant amounts of lactate produced from glucose in the *Ap*/*Lp* co-culture condition (Figure S6). To directly test if lactate production is necessary for the observed effect of the bacterial community on the behaviour of the host, we chose to genetically ablate lactate production in *Lp* and assess if this affected the ability of the bacterial community to suppress yeast appetite. We used a *Lp*^WCFS1^ strain harbouring a deletion of the *ldhD* and *ldhL* genes (*Lp*^WCFS1*ldh*^), which had been shown to be important for lactate production in this *Lp* strain (Ferain *et al.*, 1996). While the co-culture of *Ap* with the *wt Lp*^WCFS1^ control strain strongly reduced the Ile deprivation induced increase in yeast feeding, the *Lp*^WCFS1*ldh*^ mutant strain failed to suppress yeast appetite (Figure 4A). The failure of the *Lp*^WCFS1*ldh*^ mutant strain to suppress yeast appetite could be compensated by adding back lactate to the medium, confirming the conclusion that lactate production by *Lp* is necessary for the community to alter the food choice of the host. These data clearly show that lactate production by *Lp* is necessary for the commensal bacteria community to alter food choice.

**Fig. 4.**
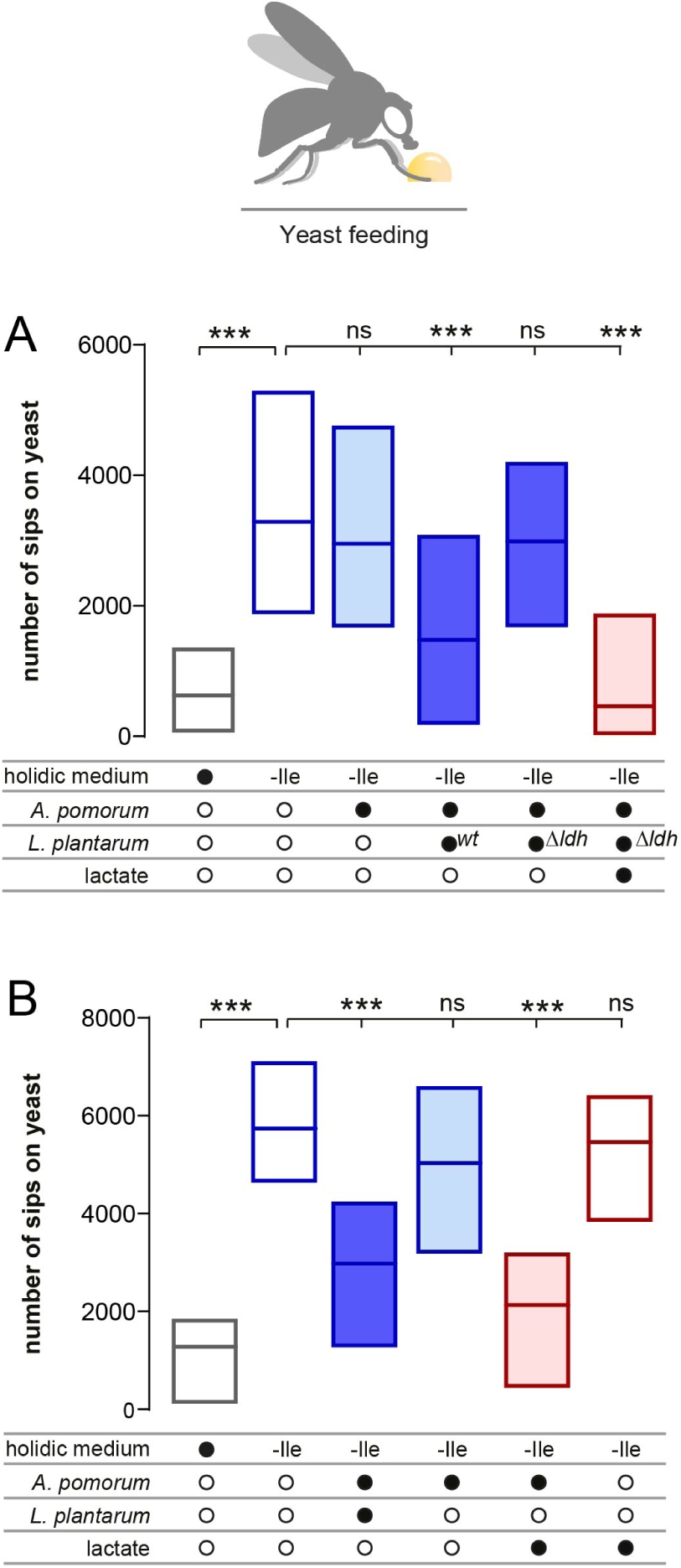
Lactate production by *Lp* is necessary and sufficient to reduce yeast appetite in the presence of *Ap*. (A and B) Number of sips on yeast of either germ-free flies (empty boxes) kept in complete holidic medium (grey border boxes) or medium without Ile (-Ile, blue border boxes) of flies that were monoassociated (light blue) with *Ap*, or biassociated (dark blue) with both *Ap* and *Lp*, or grown in medium containing lactate (bordeaux). Filled black circles represent a complete holidic medium or association with either *Ap*, wt or mutant *Lp* or the presence of lactate in the medium. In (A) wt denotes the parental *Lp*^WCFS1^ strain and *ldh* the deletion mutant *Lp*^WCFS1*ldh*^. Open black circles represent the absence of specific bacteria or lactate. Boxes represent median with upper and lower quartiles. (A) n = 51-85. (B) n= 33-38. Significance was tested using a Kruskal–Wallis test with a Dunn’s multiple comparison test. Not significant (ns) p > 0.05, *** p 0.001.

These results prompt the intriguing possibility that lactate production is the only critical metabolite provided by *Lp* for the commensal bacteria community to be able to exert its effect on host behaviour. To test this hypothesis we assessed if lactate is sufficient to replace *Lp* in its ability to affect host behaviour in the context of the commensal bacterial community. As expected, flies in which *Lp* was removed from the bacterial community showed the same Ile deprivation induced yeast appetite as germ-free animals (Figure 4B). Strikingly, replacing *Lp* with lactate in the *Ap* gnotobiotic animals lead to a potent suppression of yeast appetite, despite the host being Ile deprived for multiple days (Figure 4B). This effect was not due to a direct, unspecific effect of lactate on the host as lactate alone (with-out *Ap*) did not affect yeast appetite. These data together with the data from the lactate production mutant clearly show that lactate is the key metabolite provided by *Lp* allowing the bacterial community to alter food choice. Furthermore, these data strongly suggest that *Ap* is the main microbial player altering host behaviour and that *Lp* acts mainly as a provider of lactate, which could be required for *Ap* to synthesize the critical factors that drive the alterations in feeding behaviour.

### Lactate is used by *Ap* to produce amino acids

What could be the metabolites synthesized from lactate by *Ap* required for the bacterial community to act on the host? Given that the synthesis and secretion of AAs by *Ap* was increased by the presence of *Lp*, we wondered whether lactate could serve as a precursor for the production of these important nutrients. To test this hypothesis, we cultivated *Ap* in holidic media in which we replaced *Lp* with uniformly labelled ^13^C_3_-L-lactate (20 g/l). We tracked the synthesis of ^13^C labelled AAs from this carbon source, normally provided by *Lp* using LC-MS. Confirming our hypothesis, in the supernatant of *Ap* cultures grown in media lacking Ile, we found higher levels of the m+3, m+4 and m+5 forms of ^13^C-Ile, compared to the levels found in the no growth and no-labelling ^12^C-glucose control cultures (Figure 5). Furthermore, we could detect synthesis from lactate of all AAs that were previously detected as being produced from glucose in *Ap*/*Lp* co-cultures (Figure S7). Importantly, the same pattern of AA labelling was observed in 48h cultures with the expected exception that heavier isotopomers were also detected (Figure S8). Furthermore the AA labelling in the 24h and 48h samples showed very similar patterns as the one detected in *Ap* and *Lp* co-cultures using labelled glucose. Our results show that *Ap* synthesizes multiple AAs from lactate and are in agreement with earlier reports that lactate can serve as an important precursor for AA synthesis in other *Acetobacteraceae* (Adler *et al.*, 2014). Interestingly, the average amount of ^13^C-Ile synthesized from ^13^C_3_-lactate was similar to that measured in the co-cultures of *Ap* and *Lp* synthesized from ^13^C-glucose (Figure 3 and 5 and Figure S5, S7 and S8), enforcing the similarity between the co-culture condition and the lactate *Ap* condition. Given that our original behavioural and culture experiments included His deprivation, we also analysed the levels of *de novo* synthesized His, an eAA for the fly and a non-essential amino acid (neAA) for *Ap* and *Lp* (Figure S1 and S2). His levels were overall similar in all diets both in the *Ap* single cultures and in co-cultures in which labelled glucose was used as a precursor (Figure S4 and S5), and in *Ap* cultures where labelled lactate was used as a tracer (Figure S7 and S8). Interestingly, no His was detected in co-cultures growing in media lacking His. This suggests that *Lp* prioritizes the consumption of AAs existing in the media over its synthesis (Figure S4 and S5). Altogether, these results show that *Ap* uses lactate to synthesize secreted AAs. In the host, *Lp* derived lactate is therefore very likely to be used by *Ap* to synthesize AAs which are then used by *Lp* to grow and produce lactate, allowing the community to overcome deleterious dietary conditions. The syntrophic relation between *Ap* and *Lp* therefore ensures the dietary stability of the community, while also ensuring a constant flux of lactate to *Ap* enabling this bacterium to exert its function on host behaviour.

**Fig. 5.**
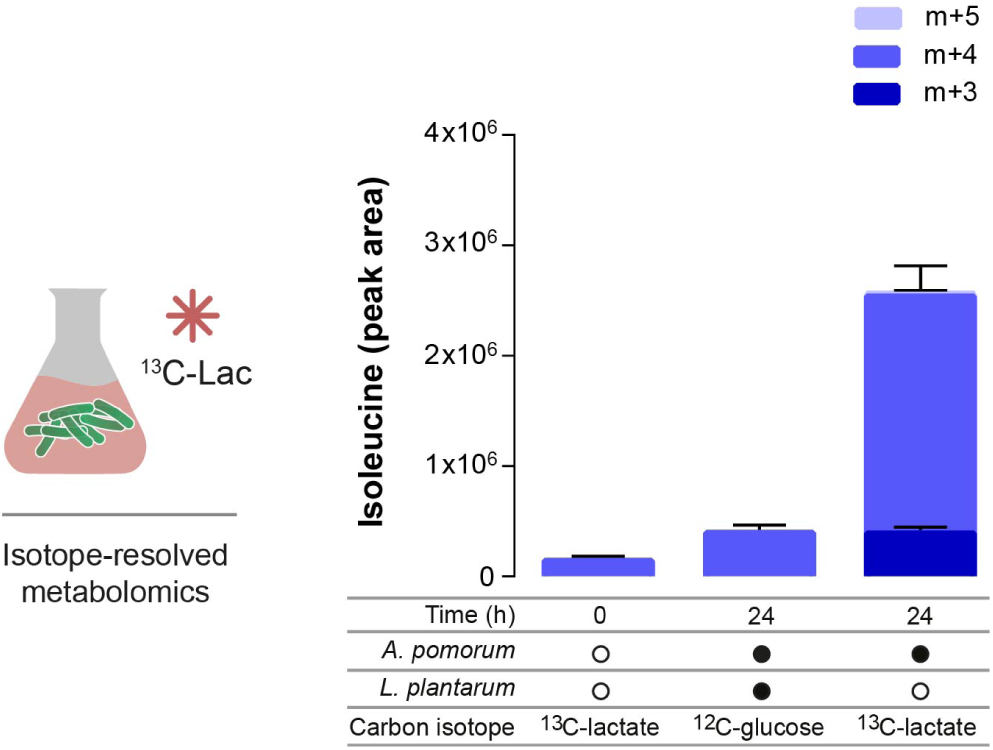
*Ap* synthesizes Ile from lactate. The stacked bars represent the relative amount of Ile isotopomers measured in the supernatant of liquid holidic medium of bacterial cultures lacking Ile and either containing, unlabelled glucose (^12^C-glucose) as a control for unspecific labelling, or uniformly ^13^C labelled lactate (^13^C-lactate, 20 g/l). Heavy labelled Ile isotopomers were measured using LC-MS in samples collected after 24h of bacterial growth and displayed as metabolite peak area. No bacteria were added in media representing time “0”. The number of heavy carbons incorporated per Ile molecule is indicated as m+n, where n = the number of ^13^C. Filled black circles represent the presence of specific bacteria in the culture. Open black circles represent the absence of specific bacteria. Each data point represents the mean of three biological replicates and the error bars represent the standard error of the mean for each isotopomer.

### AAs *de novo* synthesized by bacteria are unlikely to suppress flies’ yeast appetite

So far, our results show that *Ap* sustains *Lp* growth *in vivo* and *in vitro* in absence of Ile and that either in the presence of *Lp* or lactate, *Ap* increases the synthesis and secretion of AAs. It is widely accepted that gut microbes are an important source of eAAs in different hosts, insects and humans included (Douglas and Prosser, 1992; Metges, 2000). Our data would therefore be compatible with a simple model in which *Ap* synthesized AAs would not only be required to ensure the growth of *Lp* but would also act on the host to suppress yeast appetite. If this would be the case, the amount of AAs provided by the bacterial community should be comparable with the amount of dietary AAs sufficient to suppress protein appetite in the axenic animals. We therefore first decided to compare the amount of Ile secreted by the bacterial community with the amounts measured in the complete fly medium, which we know efficiently suppresses yeast appetite. While in the Ile dietary deprivation situation we could detect *de novo* synthesized Ile in the co-culture as well as in the *Ap* culture supplemented with lactate (Figure 3 and 5), the amount of secreted Ile in these conditions is a 1/1000^th^ of the amount present in the complete holidic medium (Figure 6A). In order to identify the amount of Ile required to suppress yeast appetite we next titrated the concentrations of Ile in the holidic medium and tested the feeding behaviour of axenic flies maintained on these diets. Strikingly, a diet with 25% of the total concentration of Ile did not lead to a significant suppression of yeast appetite, suggesting that this amount of Ile is not sufficient to significantly alter the behaviour of the fly (Figure 6B). Only the addition of at least 50% of the full Ile concentration led to a complete suppression of protein appetite. This shows that commensal bacteria would have to provide relatively high amounts of dietary Ile (between 25 and 50% of the original amounts in the holidic medium) to directly suppress yeast appetite. Our *in vitro* measurements, however, suggest that the bacterial community secretes orders of magnitude lower amounts of Ile than the ones required to suppress yeast appetite (Figure 6A). This makes it unlikely that the amount of AAs secreted by the bacterial community is sufficient to suppress protein appetite.

**Fig. 6.**
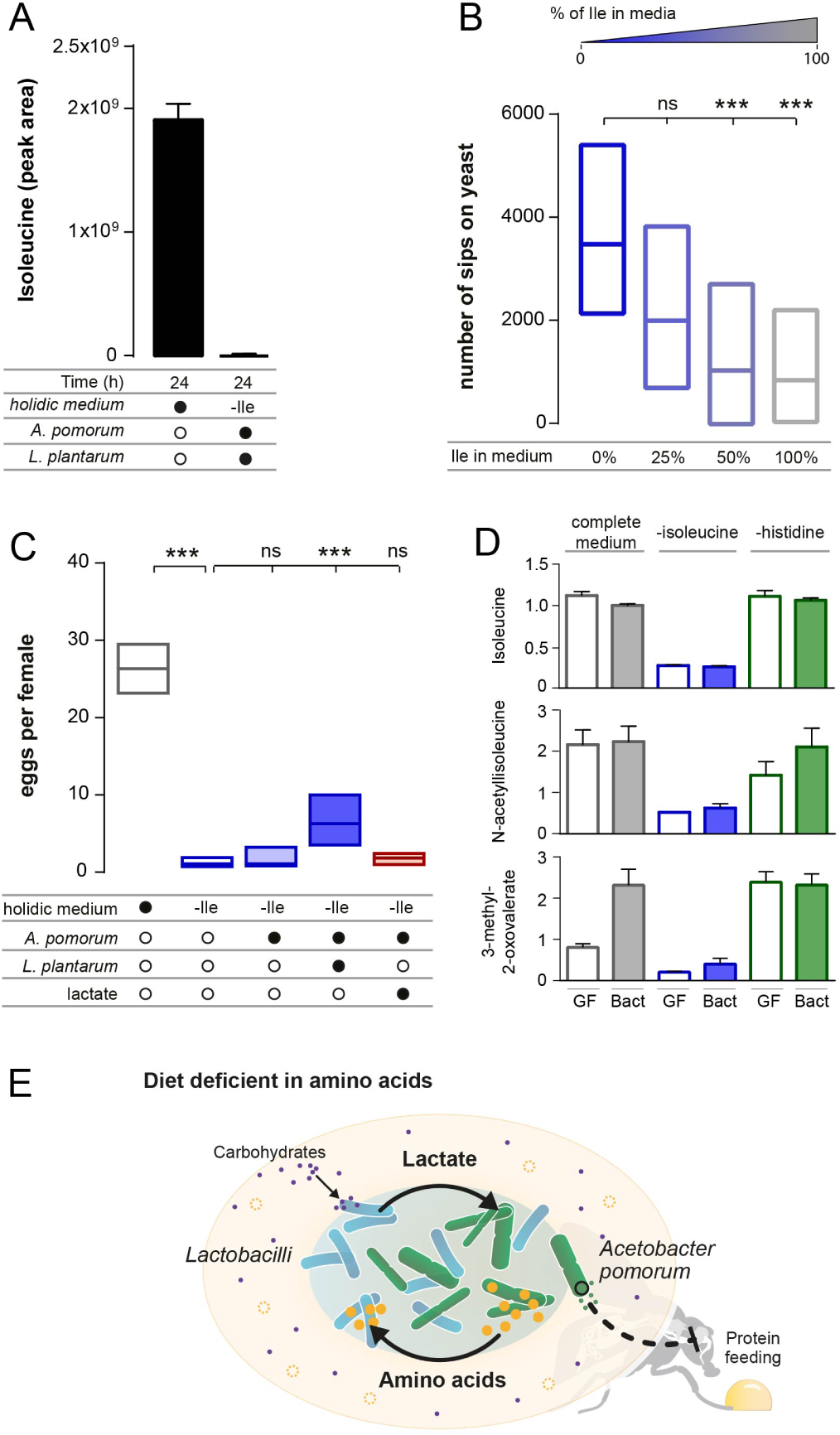
Bacterial amino acid supply are unlikely to explain the effect of gut bacteria on behaviour. (A) Relative amount of total (labelled plus unlabelled) Ile in complete holidic medium and medium lacking Ile, determined by LC-MS in sterile media or in media in which *Ap* and *Lp* were co-cultured. (B) Number of sips on yeast of flies maintained in holidic medium lacking Ile (0%), or in holidic media containing 25%, 50% or 100% of the total Ile amount found in complete medium). n = 44-53. (C) Number of eggs per mated female flies. Axenic flies were maintained on complete holidic medium or medium lacking Ile (open boxes). Flies monoassociated with *Ap* (light blue box), flies co-associated with *Ap* and *Lp* (dark blue box), and flies associated with *Ap* and maintained in media supplemented with lactate (bordeaux box), where maintained on holidic medium lacking Ile. n=23. (D) Bars represent the relative abundance of Ile and Ile degradation metabolites as measured in metabolomics experiments from heads of germ-free flies (GF) and flies inoculated with commensal bacteria (Bact) and maintained in either complete holidic medium (grey boxes), medium lacking Ile (blue boxes) or medium lacking His (green boxes). n=5. (E) Schematic describing the metabolic interactions resulting from the syntrophic relation between *Ap* and *Lp* and their potential effects on the bacterial community and the host. The carbohydrates in the host diet are represented as purple dots, AAs as orange circles and the absence of eAAs in the diet is represented as dashed circles. *Lp* and *Ap* are represented as blue and green bacteria, respectively. Black arrows represent the flow of lactate produced by *Lp* and utilized by *Ap* to produce AAs which are then used by *Lp* in situations of dietary AA scarcity. The black dashed inhibitory arrow indicates the effect of *Ap* in suppressing protein appetite of the host through a yet undetermined mechanism requiring the presence of lactate produced by *Lp*. In (A and C) filled black circles represent a complete holidic medium or association with specific bacteria or the presence of lactate. Open black circles represent the absence of specific bacteria or lactate in the diet. (A and D) Data are plotted as the mean of three (A) or five (D) biological replicates and the error bars represent the standard error of the mean. (B) Significance was tested using a Kruskal–Wallis test with a Dunn’s multiple comparison test. Boxes represent median with upper and lower quartiles. Not significant (ns) p > 0.05, *** p 0.001.

We decided to back this conclusion using a different physiological readout for AA availability. In *Drosophila* dietary eAAs are the main rate limiting nutrients required for egg production (Leitão-Gonçalves *et al.*, 2017; Piper *et al.*, 2014). As such egg laying can be used as an almost linear readout for the physiological availability of eAAs (Piper *et al.*, 2017). We had shown that commensal bacteria can increase egg laying in eAA deprived animals (Leitão-Gonçalves *et al.*, 2017). Indeed, we observed that Ile deprivation led to a drastic decrease in egg laying in germ-free females, which was mildly rescued in females harbouring an *Ap*/*Lp* community (Figure 6C). This can be interpreted as the community providing AAs to the host which it then uses for producing eggs. It is however important to note that the rate of egg laying is still very low when compared to what would be expected on a diet containing between 25%-50% of the original Ile diet (Piper *et al.*, 2014). Even more strikingly, in flies in which we replaced *Lp* by lactate, egg laying was not increased when compared to the *Ap* alone and the germ-free controls (Figure 6C). Given that providing flies with *Ap* and lactate is sufficient to completely suppress protein appetite, this shows that the effect of the commensal bacteria on behaviour can be functionally separated from the effect on egg laying. The failure of the *Ap*+lactate condition to rescue egg laying in an –Ile situation, strongly supports our earlier conclusion that the amount of Ile provided by *Ap* is not sufficient to provision the fly with adequate amounts of this essential nutrient. This strengthens the evidence that bacterially synthesized AAs are unlikely to contribute to the suppression of yeast appetite. Finally, we tested if we could detect an increase in free amino acids in AA deprived flies with a bacterial community. For this we performed metabolomics on isolated heads of germ free and gnotobiotically bacterially reconstituted females (to avoid the confounding contribution of reducing the amount of eggs by eAA deprivation). Supporting previous results (Leitão-Gonçalves *et al.*, 2017), in axenic conditions the removal of either Ile or His drastically reduced the levels of the corresponding free amino acid (Figure 6D and Figure S9). This finding is supported by the observed concomitant reduction in the levels of multiple metabolites derived from these amino acids such as N-acetylisoleucine, 3-methyl-2-oxovalerate, N-acetylhistidine and imidazole lactate (Figure 6D and Figure S9). In agreement with all our results the presence of gut bacteria did not rescue this reduction in free amino acids and corresponding derived metabolites (Figure 6D and Figure S9). The levels of the deprived eAAs and its metabolites in the fly remained very low in both the flies with and without gut bacteria. These measurements made in the host further support the conclusion that gut bacteria are unlikely to act on host behaviour by rescuing the levels of free AAs in the animal. Overall our data are in agreement with a model in which *Lp* and *Ap* grow as a community in the fly where they engage in a syntrophic interaction buffering them from adverse dietary host conditions (Figure 6E). *Lp* provides lactate to *Ap* which it uses to synthesize and secrete AAs. This ensures *Lp* growth even in detrimental dietary conditions in which limiting AAs are missing. This also ensures the constant flux of lactate which provides the necessary fuel for *Ap* to synthesize the metabolites that alter choice behaviour (Figure 6E) which according to our data are unlikely to be proteogenic AAs. Syntrophic relations between gut bacteria could therefore be a common theme among gut bacterial communities, generating metabolic cycles which buffer them from suboptimal host dietary conditions and allowing them to generate a constant flux of metabolites acting on the host.

## Discussion

Identifying the mechanisms by which metabolic exchanges shape diet-microbiome-host interactions is key to understanding how gut microorganisms alter the physiology and behaviour of the host. Both the food choice of animals and their microbiome are altered by changes in diet. How one mechanistically relates to the other is currently poorly understood. We had shown that two bacteria abundant in fly’s microbiome (*Ap* and *Lp*) act together to suppress the protein appetite of AA deprived animals (Leitão-Gonçalves *et al.*, 2017). Here we show that they need to act as a community to establish a syntrophic relationship that enables them to overcome drastic nutritional limitations generated by imbalanced host diets. While most studies on the impact of host diet on the microbiome emphasize the ability of diet to alter the microbiome (David *et al.*, 2014; Johnson *et al.*, 2019; Turnbaugh *et al.*, 2009; Wang *et al.*, 2014) our work highlights a different facet of diet-microbiome relationships: how metabolic interactions within the microbiome allow the gut microbiome to become resilient to changes in the host diet. Key is the ability of multiple microbes to establish communities in which syntrophic relationships allow specific members of the community to overcome auxotrophies for specific nutrients. While widely studied in the context of microbial ecology (Mee *et al.*, 2014; Morris *et al.*, 2013; Ponomarova and Patil, 2015) less is known about the metabolic interactions shaping gut microbes and their importance in how they act on the host. This is especially the case in humans where we have just started analysing the nutritional preferences and metabolic idiosyncrasies of single members of the human gut microbiome (Tramontano *et al.*, 2018). While it has been proposed that microbial metabolic interactions might be key for the generation of specific effector metabolites such as GABA or serotonin (Olson *et al.*, 2018; Sharon *et al.*, 2014), we show that an important aspect of gut microbial communities is their resilience towards dietary perturbations, thereby ensuring a stable and constant impact of the microbiome on the host.

Lactic acid bacteria are known to often co-occur with other microorganisms in a variety of natural niches (Duar *et al.*, 2017; Ponomarova and Patil, 2015). One such niche is the adult *Drosophila* gut where *Acetobacteracaea* and *Lactobacilli* are often found together (Pais *et al.*, 2018; Ren *et al.*, 2007; Ryu *et al.*, 2008; Wong *et al.*, 2011). The co-occurrence of bacteria from these two genera in the fly has been shown to increase the propagation of both bacteria and to contribute to the impact of the microbiome on physiological traits of the host (Gould *et al.*, 2018; Newell and Douglas, 2014; Sommer and Newell, 2019). Although lactate has been proposed to play a role in these interactions (Sommer and Newell, 2019) the mechanisms that promote and sustain bacterial growth in co-cultures and how they modify the host remain largely unexplored. Lactobacilli lack many key genes required for the synthesis of different essential nutrients such as AAs and vitamins (Martino *et al.*, 2016; Newell *et al.*, 2014; Wu *et al.*, 2017). Conversely their genome encodes an unusually large repertoire of transporters highlighting their ability if not requirement to take up nutrients from their environment (Kim *et al.*, 2013; Martino *et al.*, 2016). Using a chemically defined diet (Piper *et al.*, 2017), we show that *Lp*, a bacteria auxotrophic for Ile, is able to proliferate *in vitro via* the uptake of Ile produced by *Ap*. Moreover, the cross-feeding of this eAA, most likely, occurs also *in vivo*, since *Lp* levels are higher in biassociated than in monoassociated flies deprived of Ile and *Lp* is necessary with *Ap* for altering feeding behaviour and egg laying in these flies. Microbial cross-feeding has been shown to allow intra and inter-species exchange of several nutrients, including AAs, through its secretion to the media or *via* bacterial nanotubes (Mee *et al.*, 2014; Shitut *et al.*, 2019; Ziesack *et al.*, 2019). The *Ap*-*Lp* interaction is however not unidirectional as has been shown for example for yeast-*Lactobacilli* interactions (Ponomarova *et al.*, 2017). The production and secretion of AAs by *Ap* depends or is strongly enhanced by the presence of *Lp*. Concomitantly, this interaction also spurs the growth of *Ap* as this bacterium grows better in the fly in the presence of *Lp*. This positive effect of *Lp* on AA synthesis by *Ap* is best explained by the preferential use of lactate by *Ap* for the production of AAs (Adler *et al.*, 2014) which coincidentally is one of the main metabolic by-products produced and secreted by *Lp*. These metabolic interactions within the *Ap*/*Lp* community allow these two bacteria to create a “circular economy” in which they both optimally use the available nutritional resources provided by the host diet, allowing them both to overcome detrimental host diets and boosting their metabolic output.

Our results also strongly suggest that one or multiple metabolites derived from *Ap* lactate metabolism are likely to be the effectors altering feeding behaviour in flies co-inoculated with *Ap* and *Lp*. This conclusion is based on the observation that lactate can fully substitute *Lp* to suppress yeast appetite in *Ap*-monoassociated flies deprived of Ile. One straightforward hypothesis is that *Ap*, an autotrophic strain, synthesizes and provides eAAs to the host, suppressing protein appetite. In fact, there is evidence that gut microbes can supply significant amounts of essential nutrients, like AAs, to the host including humans (Douglas and Prosser, 1992; Metges, 2000; Sannino *et al.*, 2018; Smith *et al.*, 2013). However, we find that the bacteria produce and secrete extremely low amounts of AAs compared to those existing in the complete holidic medium which has been optimized to promote egg laying and lifespan of the fly (three orders of magnitude less). This significantly weakens the hypothesis that *Ap* alters yeast appetite by supplying enough eAA to the host so that it can compensate for the dietarily absent eAA. Especially considering that our psychometric measurements indicate that concentrations of 50% or more of the Ile concentrations in the media are necessary to significantly suppress protein appetite. This strongly suggests that the gut bacteria would need to produce at least that amount of AAs to alter protein appetite *via* these metabolites, which is not compatible with our *in vitro* and *in vivo* measurements. Moreover, we have shown that *Ap* and *Lp* are required together to supress yeast appetite in His-deprived flies (Leitão-Gonçalves *et al.*, 2017). Multiple evidence however suggest that His production by the bacterial community does not explain the suppression in yeast appetite observed in His-deprived flies: our *in vitro* measurements show that 1) while *Ap* is not sufficient to suppress yeast appetite on its own it produces and secretes His when cultivated in isolation (Figure S4) and that 2) while in a –His diet the *Ap*/*Lp* co-culture is able to suppress protein appetite, we can hardly detect His synthesis and secretion, suggesting that synthesized His is exhausted from the media by the bacterial community when it is dietarily absent. These data make it further unlikely that the bacteria reduce yeast appetite of the fly by rescuing the absence of His in the diet. Finally, the mono-association of malnourished flies with *Ap* in the presence of lactate does not increase egg laying, a physiological process which is profoundly dependent on AA availability and we could not detect a rescue in the heads of flies associated with bacteria of the free AAs which had been removed from the diet. All these results together with earlier published data (Leitão-Gonçalves *et al.*, 2017), fail to support the straight forward hypothesis that the bacteria act on yeast appetite by producing enough eAAs to replace the effect of the dietary removal of specific eAAs.

It is however important to note that in the physiological *Lp*/*Ap* bias-sociation situation, the presence of the microbiome is beneficial for the animal as it allows the fly to lay more eggs in an eAA deprived diet situation. This might sound counterintuitive to the above mentioned arguments as egg laying requires building blocks. But this apparent contradiction can be easily resolved by our finding that egg laying and protein appetite can be functionally separated. The modest improvement in egg laying could simply result from the use of the increased bacterial biomass in the biassociation situation, which would be sufficient to support a modest amount of egg laying while not being sufficient to suppress yeast appetite. This interpretation is supported by our finding that in the *Ap*+lactate situation protein appetite is completely abolished while egg laying is not increased. In a malnutrition setting the *Ap*/*Lp* community is hence beneficial for the fly as it allows it to maintain egg laying. Importantly, this benefit requires the ability of the community to grow in a diet lacking AAs for which *Lp* is auxotrophic. The ability of the bacterial community to withstand dietary perturbations is therefore key for its beneficial effect on the adult host.

If the bacterial community does not act on food choice behaviour by maintaining a high level of eAAs in the host, how does it then modify behaviour? The here presented data support the hypothesis that it is one or multiple, lactate-derived, *Ap*-generated metabolite(s) which modulate the feeding behaviour of the fly. This fits with earlier data suggesting that the gut bacteria need to be metabolically active to modify food choice (Leitão-Gonçalves *et al.*, 2017). Gut bacteria can contribute with a plethora of small metabolites on concentrations comparable to those administered in drug doses (10 µM–1 mM) (Nicholson *et al.*, 2012). These include neuroactive substances such as GABA and serotonin (Strandwitz *et al.*, 2019; Yano *et al.*, 2015). Furthermore, recent studies identified multiple bacterial metabolites acting as G-protein-coupled receptors (GPCRs) agonists, which have the potential to affect host physiology, including the nervous system (Chen *et al.*, 2019; Colosimo *et al.*, 2019). The reduction of the complexity of the behaviourally active community to one bacterium (*Ap*) and the identification of lactate as a likely precursor for the generation of the neuroactive metabolites allows a targeted focus to identify the precise mechanisms by which the gut microbiome alters food choice. A combination of metabolomics approaches, including stable isotope-resolved metabolomics, and bacterial genetics, including unbiased genetic screens, should allow for the identification of the exact molecular mechanisms underlying the changes in behaviour.

Our work identifies the mechanisms that sustain a syntrophic relation between two abundant species of bacteria found in the fly’s microbiome and identifies the species that drives the alteration in behaviour observed in the host. We also show that this mutualistic relationship buffers the effect of dietary restrictions in both, the microbiome and the host. Diet is an essential, dynamic, and highly diverse environmental variable deeply affecting several aspects of behaviour, including food choice (Leitão-Gonçalves *et al.*, 2017; Simpson *et al.*, 2015; Solon-Biet *et al.*, 2019; Tarlungeanu *et al.*, 2016), as well as the microbiome, which has been shown to play a critical role in human behaviour, including through the metabolism of nutrients and drugs (Dodd *et al.*, 2017; Kessel *et al.*, 2019; Rekdal *et al.*, 2019; Sharon *et al.*, 2019; Smith *et al.*, 2013). Our study highlights the importance of metabolic interactions among different species of gut bacteria in shaping the outcome of diet on the microbiome and its impact on host physiology and behaviour. Given the numerical complexity of the human microbiome and its high inter-individual heterogeneity (The Human Microbiome Project Consortium *et al.*, 2012) mechanistically disentangling such relationships is a daunting task. The ability to combine extremely precise dietary manipulations using a holidic diet, as well as microbial, genetic and molecular perturbations with detailed behavioural, physiological and metabolic phenotyping in high throughput makes *Drosophila* an ideal system to identify mechanisms by which gut bacterial communities act on the host. Importantly, given that the key molecular and physiological regulatory mechanisms are conserved across phyla, the microbiome mechanisms identified in invertebrates have been shown to be translatable to vertebrates (Schwarzer *et al.*, 2016). Our study shows the potential in using *Drosophila* to mechanistically disentangle the influence of diet on microbiological communities and identify the individual contributions of bacterial species on host behaviour and brain function. The identified molecular and metabolic strategies can then be harnessed to explore similar mechanisms in vertebrates, including humans, providing an attractive path for the efficient mechanistic dissection of how gut microbes act on the host across phyla.

## Acknowledgements

We thank the laboratory of François Leulier (IGFL, France) for providing bacterial strains and Jessika Consuegra for advice on the proper cultivation conditions for bacterial strains and suggestions on how to test the effect of lactate on the host. We thank Matthew Piper, Gili Ezra, Ibrahim Tastekin, Darshan Bharatkumar Dhakan, Daniel Münch, and members of the Behaviour and Metabolism laboratory for helpful discussions and comments on the manuscript, and to Gil Costa for illustrations. We thank the Glass wash and media preparation and the fly platforms at the Champalimaud Centre for the Unknown for technical assistance. This project was supported by funding from the Kavli Foundation to C.R.. The project leading to these results has received funding from “la Caixa” Banking Foundation to C.R. under the project code HR-17-00539. A.P.F. is a member of the Champalimaud International Neuroscience Doctoral Programme and is supported by the FCT fellowship PD/BD/114277/2016. This project was supported by the Portuguese Foundation for Science and Technology (FCT) postdoctoral fellowship SFRH/BPD/79325/2011 to Z.C-S. Work by Z.C-S. was also financed by national funds through the FCT, in the framework of the financing of the Norma Transitória (DL 57/2016). O.D.K.M. is supported by the CRUK Career Development Fellowship, C53309/A19702. Research at the Centre for the Unknown is supported by the Champalimaud Foundation.

## Author contributions

Conceptualization, S.F.H. and C.R.; Data Curation and formal analysis, S.F.H., L. S., Z. C., O. D. K. M., C.R.; Investigation, S.F.H. and L. S. except from: metabolomics analysis by LC-MS which was conducted by T. Z. and acquisition of samples for metabolomics analysis in the heads of flies which was conducted by A. P. F., Z. C., B., A. P. E. and M. A.; Supervision, C. R.; Validation, S.F.H. and C.R.; Visualization, S.F.H, L.S. and C.R.; Writing – Original Draft, S.F.H. and C.R.; Writing – Review Editing, S.F.H., L.S. and C.R.; Project Administration, C.R.; Funding acquisition, C.R.. All authors read and approved the final manuscript.

## Materials and Methods

### *D. melanogaster* husbandry and dietary treatments

All experiments were performed with either axenic or gnotobiotic mated w^1118^ female flies. Flies were reared under controlled conditions at 25°C, 70% humidity, and 12 h light/dark cycle. Axenic fly stocks were generated as described in ((Leitão-Gonçalves *et al.*, 2017) and dx.doi.org/10.17504/protocols.io.hebb3an) and maintained on high yeast-based medium (highYBM) (per litre of water: 8 g agar [NZYTech, PT], 80 g barley malt syrup [Próvida, PT], 22 g sugar beet syrup [Grafschafter, DE], 80 g corn flour [Próvida, PT], 10 g soya flour [A. Centazi, PT], 60 g instant yeast [Saf-instant, Lesaffre], 8 ml propionic acid [Argos], and 12 ml nipagin [Tegospet, Dutscher, UK] [15% in 96% ethanol]) laced with antibiotics (50 µg/ml tetracycline, 50 µg/ml kanamycin, 50 µg/ml ampicillin, and 10 µg/ml erythromycin). The absence of bacteria was assessed regularly by grinding flies in sterile 1x PBS and spreading the suspension on LB, MRS, or Mannitol plates. For the experiments described in this study axenic fly cultures were set in sterile highYBM without antibiotics using 6 females and 4 males per vial, to guarantee a homogeneous density of offspring across the different experiments, and left to develop until adulthood in this media. Holidic medium (HM) without preservatives and with an optimizes AA composition (FLYAA) was prepared as described in ((Leitão-Gonçalves *et al.*, 2017), and dx.doi.org/10.17504/protocols.io.heub3ew). HM lacking isoleucine or histidine was prepared by simply omitting the corresponding AAs. A 300 g/l lactic acid solution (pH 4.0-4.5) was prepared from a 90% (w/w in H2O) DL-lactic acid solution (Sigma-Aldrich, 69785) using 10 M NaOH, and used to prepare HM supplemented with lactate at 20 g/l. Flies were exposed to the different dietary treatments using the protocol described in ((Leitão-Gonçalves *et al.*, 2017) and dx.doi.org/10.17504/protocols.io.hhcb32w) where axenic or gnotobiotic 1 to 5-d-old flies were used. Briefly, 16 females and 5 males were placed in highYBM-containing vials (without preservatives), transferred to fresh highYBM after 48h and transferred, after 24h, to the different HM (sterile or bacteria-treated media) where they were maintained for 72 h before any indicated assay.

### Bacterial species and generation of pre-inocula

The following bacteria strains were used in this study: *Lactobacillus plantarum*^NC8^ (Axelsson *et al.*, 2012), *Lactobacillus plantarum*^WCFS1^ (Ferain *et al.*, 1996), *Lactobacillus plantarum*^WCFS1*ldh*^ (Ferain *et al.*, 1996) and *Acetobacter pomorum* (Ryu *et al.*, 2008). To generate the 5 bacteria community used for the fly head metabolomics measurements the following bacterial strains were also used: *Lactobacillus plantarum*^WJL^ (Ryu *et al.*, 2008), *Lactobacillus brevis*^EW^ (Ryu *et al.*, 2008), *Commensalibacter intestini*^A911T^ (Ryu *et al.*, 2008), and *Enterococcus faecalis* (Cox and Gilmore, 2007). All *Lactobacillus* species were cultivated in 10 ml MRS broth (Fluka, 38944) as static cultures in 14 ml culture tubes (Thermo Fisher Scientific, 150268) at 37°C for 24 h. *A. pomorum* and *C. intestini*^A911T^ were cultivated in mannitol (3 g/l Bacto peptone [Difco, 0118–17], 5 g/l yeast extract [Difco, 212750], 25 g/l D-mannitol [Sigma-Aldrich, M1902]) at 30°C for 48 h with orbital agitation (180 rev.min-1). *C. intestini*^A911T^ was cultured in 20 ml of medium in 50-ml tubes (Falcon), while *A. pomorum* was cultured in 200 ml of medium in 500-ml flasks. *E. faecalis* was cultured in 200 ml of liquid LB medium (Sigma-Aldrich, L3022) in 500-ml flasks at 37°C for 24 h at 220 rev/min.

### Generation of axenic and gnotobiotic flies

Axenic flies were generated as previously described (Leitão-Gonçalves *et al.*, 2017). To prepare mono- and biassociated flies with *A. pomorum* and *Lactobacilli* strains, vials containing HM media were inoculated with the desired bacterial species before flies were transferred. The number of CFU added per vial was as follows: *L. plantarum*^NC8^, 7.2 x 107; *L. plantarum*^WCFS1^, 5.0 x 107; *L. plantarum*^WCFS1*ldh*^, 5.6 × 107; and *A. pomorum*, 9.5 × 107. In the particular case of the gnotobiotic flies used to measure the metabolites in the fly’s head the bacterial community consisted of: *L. plantarum*^WJL^(6.4 × 107 CFU), *L. brevis*^EW^ (5.31 × 106 CFU), *C. intestini*^A911T^ (9.04 × 107 CFU), *A. pomorum* (9.5 × 107 CFU) and *E. faecalis* (1.11 × 108 CFU). The volume of culture containing the appropriate number of CFUs was centrifuged (5,000 × g, 10 min), the pellet washed three times with 1xPBS and the cells resuspended in 50 µl of 1xPBS. The same volume of sterile bacterial media was centrifuged and the PBS resulting from washing the tube three times was used as a control. Fifty microliters of cell suspension were added to each fly culture vial, and those allowed to dry for 2h before transferring flies.

### Assessment of bacterial CFUs

The selection and assessment of the number of *A. pomorum* or *L. plantarum*^NC8^ CFUs in flies co-inoculated with two bacterial species or in liquid co-cultures was done by plating samples in MRS media either supplemented with Kanamycin (50 µg/l) or Ampicillin (10 µg/l) for the selection of *L. plantarum*^NC8^ or *A. pomorum*, respectively. MRS plates supplemented with Kanamycin were incubated at 37°C, while plates supplemented with Ampicillin were incubated at 30°C. Samples of monoassociated flies were plated in MRS and incubated at 30°C or 37°C for *A. pomorum* or *L. plantarum*^NC8^, respectively. All samples were plated using an easySpiral® (InterScience, 412000) plater and the number of CFU counted with an automatic colony counter Scan® 500 (Interscience, 436000).

### Assessment of the bacterial load of flies

Flies were washed in 70% ethanol, to remove bacteria adhering to the fly cuticle, and further washed twice with sterile 1x PBS. The flies were homogenized using 50 µl of 1x PBS per fly and the homogenates serially diluted and plated in an appropriate media for the selection of the different bacteria. The number of CFU per fly was measured in at least three independent biological samples of 8 flies.

### flyPAD assays

Food choice experiments were performed using the flyPAD as described in (Itskov *et al.*, 2014). The food preferences of single gnotobiotic flies with, maintained in complete HM or deprived of single AAs, were tested in an arena with food patches containing 1% agarose mixed with either 10% yeast or 20 mM sucrose. The flies were allowed to feed for 1 h and the number of sips per animal was calculated using the previously described flyPAD analysis algorithms in (Itskov *et al.*, 2014). Non-eating flies (those flies having less than two activity bouts per assay) were excluded from the analysis. To test if glucose (100 mM) can replace sucrose (50 mM) as a carbon source in the HM, the feeding behaviour of axenic and biassociated flies was tested using the flyPAD. Axenic flies fed the glucose-based HM diet exhibited the same increase in yeast appetite when deprived of isoleucine (Figure S10). Furthermore, isoleucine deprived animals associated with *Ap* and *Lp* exhibited a decreased yeast appetite compared to the axenic animals. These data show that glucose and sucrose are interchangeable as carbon sources in what concerns the feeding behaviour of the flies.

### Egg-laying assays

After 72 h on HM with different AA compositions, groups of 16 females and 5 males were placed in apple juice agar plates (250 ml/l apple juice, 19.5 g/l agar, 20 g/l sugar, and 10 ml/l nipagin [15% in ethanol]) and allowed to lay eggs for 24 h. Living adult flies and eggs were then counted and the number of eggs was normalized by the number of living females. Data were pooled from three experiments performed independently on different days.

### Assessment of bacterial growth in holidic media

A liquid version of the HM was used to cultivate bacteria *in vitro*. The media was prepared by removing agar and cholesterol from the HM recipe to avoid turbidity in the media and replacing sucrose with 100 mM of glucose. An appropriate volume of cells was calculated in order to inoculate 20 ml of liquid HM with an initial optical density measured at 600 nm (OD_600_) of 0.05 of each bacterium. The calculated volume of bacterial culture was centrifuged (5,000 x g, 10 min), washed once with 1x PBS and the pellet resuspended in 50 µl of 1x PBS. The cell suspension was used to inoculate 20 ml of liquid HM in 100 ml Erlenmeyer flasks, which were incubated at 25 °C with orbital agitation (180 rev/min). The number of CFU in the liquid cultures was assessed 0, 1, 2, and 3 days after the growth was resumed by plating and counting colonies as described above. The bacterial growth was determined for all tested conditions in at least three independent experiments.

### Metabolomics analysis of fly heads

At least a thousand mated gnotobiotic females maintained on each dietary condition were collected after brief CO_2_ anaesthesia and snap frozen in liquid nitrogen. To separate heads from the body and collect them, the frozen flies were vortexed in Eppendorf tubes and sieved through a 710-mm and 425-mm mesh (Retsch GmbH). All the material used to handle the body parts was continuously cooled in liquid nitrogen throughout the process, to ensure that heads were kept frozen. At least 1000 heads were sent for metabolomics profiling as a paid service at Metabolon Inc, USA. The plotted relative amount of metabolites detected in the analysis was normalized according to the number of heads in each sample.

### Metabolomic analysis by liquid chromatography-mass spectrometry (LC-MS)

To trace the synthesis of the different metabolites produced by bacteria, two universally isotopically-labelled carbon sources were used: ^13^C_6_-D-glucose (Cambridge Isotope Laboratories, CLM-1396) or ^13^C_3_-lactate (Cambridge Isotope Laboratories, CLM-1579). In experiments using ^13^C_6_-D-glucose, equimolar amounts of the heavy glucose were used to completely substitute the glucose in the HM. In experiments using ^13^C_3_-lactate, 20 g/l of the heavy lactate were added to the HM formulation which normally does not contain lactate. A control culture with HM without isoleucine with unlabelled glucose was used in parallel. The liquid version of HM without cholesterol was used. To set up the cultures, the bacterial suspensions were centrifuged (5,000 x g, 10 min) and washed, and the pellet resuspended in 50 µl of 1x PBS. Five millilitres of HM were inoculated with an initial OD_600_ of 0.05 in 25 ml Erlenmeyer flasks, and incubated at 25°C with orbital agitation (180 rev/min). A sample of the supernatant was collected after 24h of incubation. For that, 50 µl of culture were centrifuged at 5,000 x g for 3 min at 4°C and 10 µl of the supernatant immediately added to 500 µl of extraction buffer (30% methanol [Merck, 1.06035], 50% acetonitrile [Sigma-Aldrich, 900667] in miliQ water). Samples were kept at −80°C until analysis. Before collecting the sample the OD_600_ of the cultures were measured to later normalize the values of the ^13^C-labelled metabolites by the growth rate of the bacteria. Liquid chromatography-mass spectrometry (LC-MS) was performed as described previously (Maddocks *et al.*, 2017): The supernatant samples were analysed on a LC–MS platform consisting of an Accela 600 LC system and an Exactive mass spectrometer (Thermo Scientific), using a ZIC-HILIC column (4.6mm×150mm, 3.5µm) (Merck) with the mobile phase mixed by A=water with 0.1% formic acid (v/v) and B=acetonitrile with 0.1% formic acid. A gradient program starting at 20% of A and linearly increasing to 80% at 30min was used followed by washing and re-equilibration steps. The total run time was 46min. The LC stream was desolvated and ionized in the HESI probe. The Exactive mass spectrometer was operated in full-scan mode over a mass range of 75–1,000m/z at a resolution of 50,000 with polarity switching. LC-MS raw data was converted was analysed by LCquan (Thermo Scientific) and MZMine 2.10 for metabolite identification and quantification. Data represents metabolite peak area after scaling for the growth of the bacteria in culture. This was done by multiplying the peak area with the ratio of the highest OD_600_ measured in all the cultures in the metabolomics experiments and the OD _600_ of the culture from which the metabolomics measurement were done. The metabolites were measured in three independent experiments.

