## Supplementary Material for "Metabolic cross-feeding allows a gut microbial community to overcome detrimental diets and alter host behaviour"

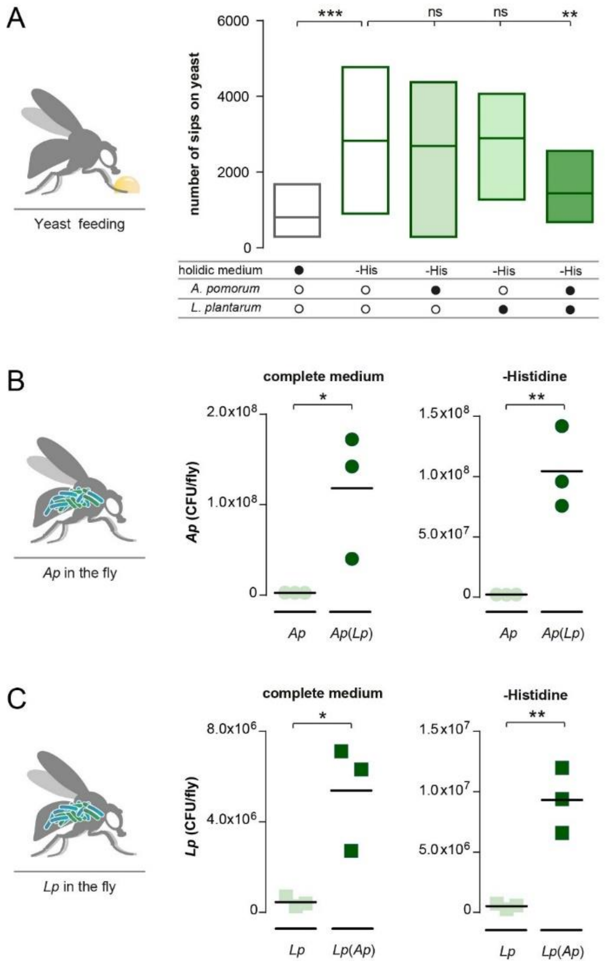


**Supplementary Figure 1 – *Ap* and *Lp* biassociation reduces yeast appetite and promotes gut bacterial growth independent of dietary Histidine content.** (A) Number of sips on yeast of flies kept in complete holidic medium or medium without His (His) that were monoassociated (light green) with *Ap* or *Lp*, or biassociated (green) with both commensal species. Boxes represent median with upper and lower quartiles. n = 51-64. Significance was tested using a Kruskal–Wallis test with a Dunn's multiple comparison test. Filled black circles represent a complete holidic medium or association with specific bacteria. Open black circles represent the absence of specific bacteria. (B and C) Number of *Ap* (circles, B) and *Lp* (squares, C) colony forming units (CFUs) from extracts of monoassociated (light blue) and biassociated (dark blue) flies kept in complete holidic media or medium lacking His (-Histidine). *Ap* and *Lp* labels data points representing the number of CFUs detected in monoassociated flies and *Ap*(*Lp*) and *Lp*(*Ap*) the measurement of *Ap* or *Lp* in biassociated flies. The black line represents the mean. n=3. Significance was tested using an unpaired t-test. Not significant (ns) $p > 0.05$, * $p \leq 0.05$, ** $p \leq 0.01$, *** $p \leq 0.001$. Given that the bacterial load experiments were all assessed in the same experiment, data plotted for the "complete medium" condition in Figure 1B and 1C and in Figure S1B and S1C, respectively, are the same.

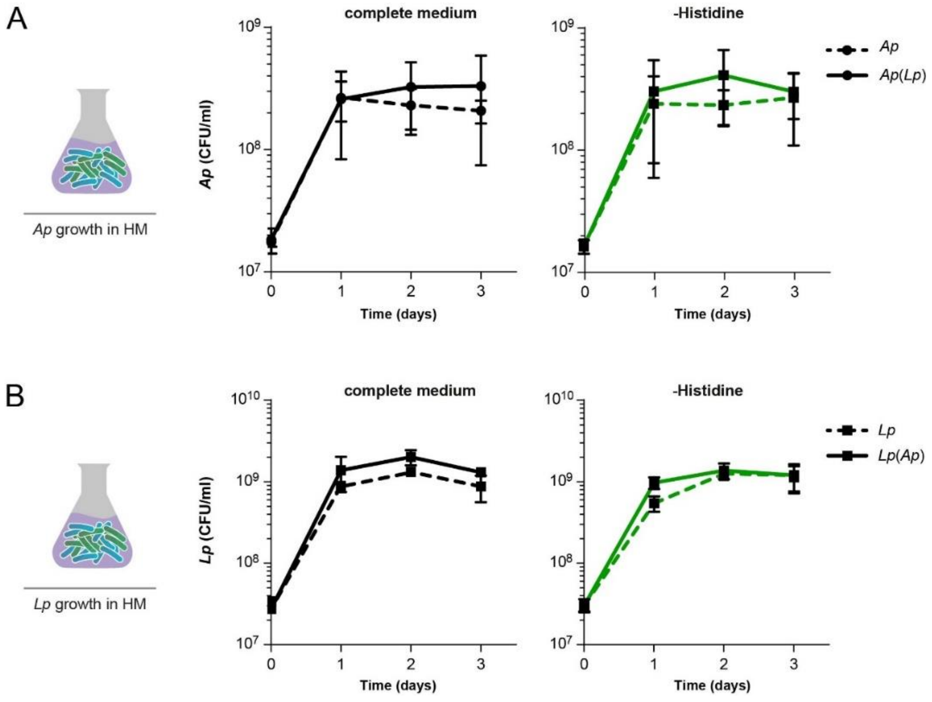


**Supplementary Figure 2 – *In vitro* growth measurements show that removing Histidine from the diet does not affect *Ap* or *Lp* growth.** *In vitro* growth curves of *Ap* (A) and *Lp* (B) in complete liquid holidic medium (black line) or in liquid medium without His (green line) plotted as CFUs at different time intervals. *Ap* and *Lp* (dashed lines) are data points representing the number of CFUs detected in monoassociated flies and *Ap*(*Lp*) and Lp(Ap) (continuous lines) the measurement of *Ap* or *Lp* in co-culture conditions. The number of CFUs per ml of culture was determined by collecting and cultivating liquid culture samples in selective media at the indicated time points (as described in material and methods). Data points represent mean values of three to four biological replicates and error bars the standard deviation of the mean. Given that number of bacteria in the *in vitro* liquid cultures were all assessed in the same experiment, data plotted for the "complete medium" condition in Figure 2A and 2B and in Figure S2A and S2B, respectively, are the same.

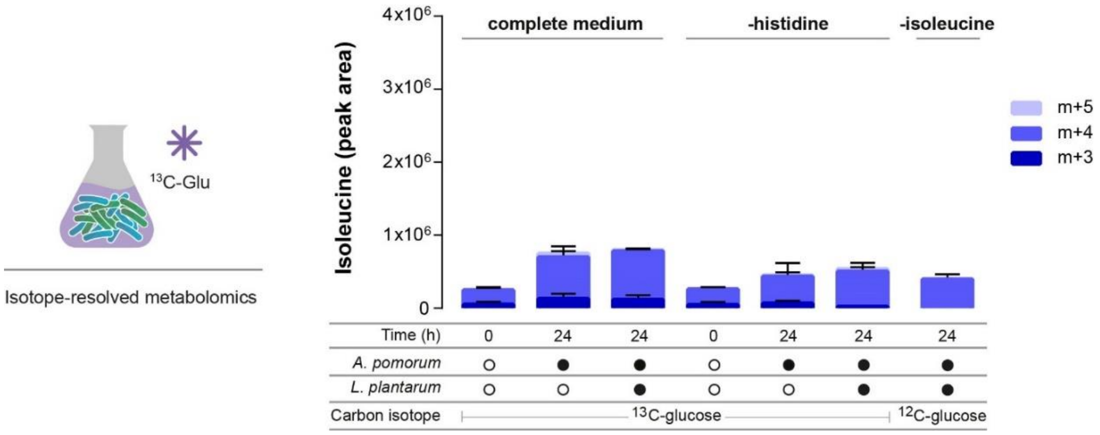


**Supplementary Figure 3 – Ile synthesis is not increased in a complete diet or in absence of His.** The stacked bars represent the relative amount of Ile isotopomers measured in the supernatant of bacterial cultures grown in complete liquid holidic medium or in medium lacking His containing uniformly $^{13}$C labelled glucose ($^{13}$C-glucose). Co-cultures were also grown in media lacking Ile and containing, unlabelled glucose ($^{12}$C-glucose) as a control for unspecific labelling. Heavy labelled Ile isotopomers were measured using LC-MS in samples collected after 24h of bacterial growth and displayed as metabolite peak area. No bacteria were added in media representing time “0”. The number of heavy carbons incorporated per Ile molecule is indicated as m+n, where n = the number of $^{13}$C. Filled black circles represent the presence of specific bacteria in the culture. Open black circles represent the absence of specific bacteria. Each data point represents the mean of three biological replicates and the error bars represent the standard error of the mean for each isotopomer.

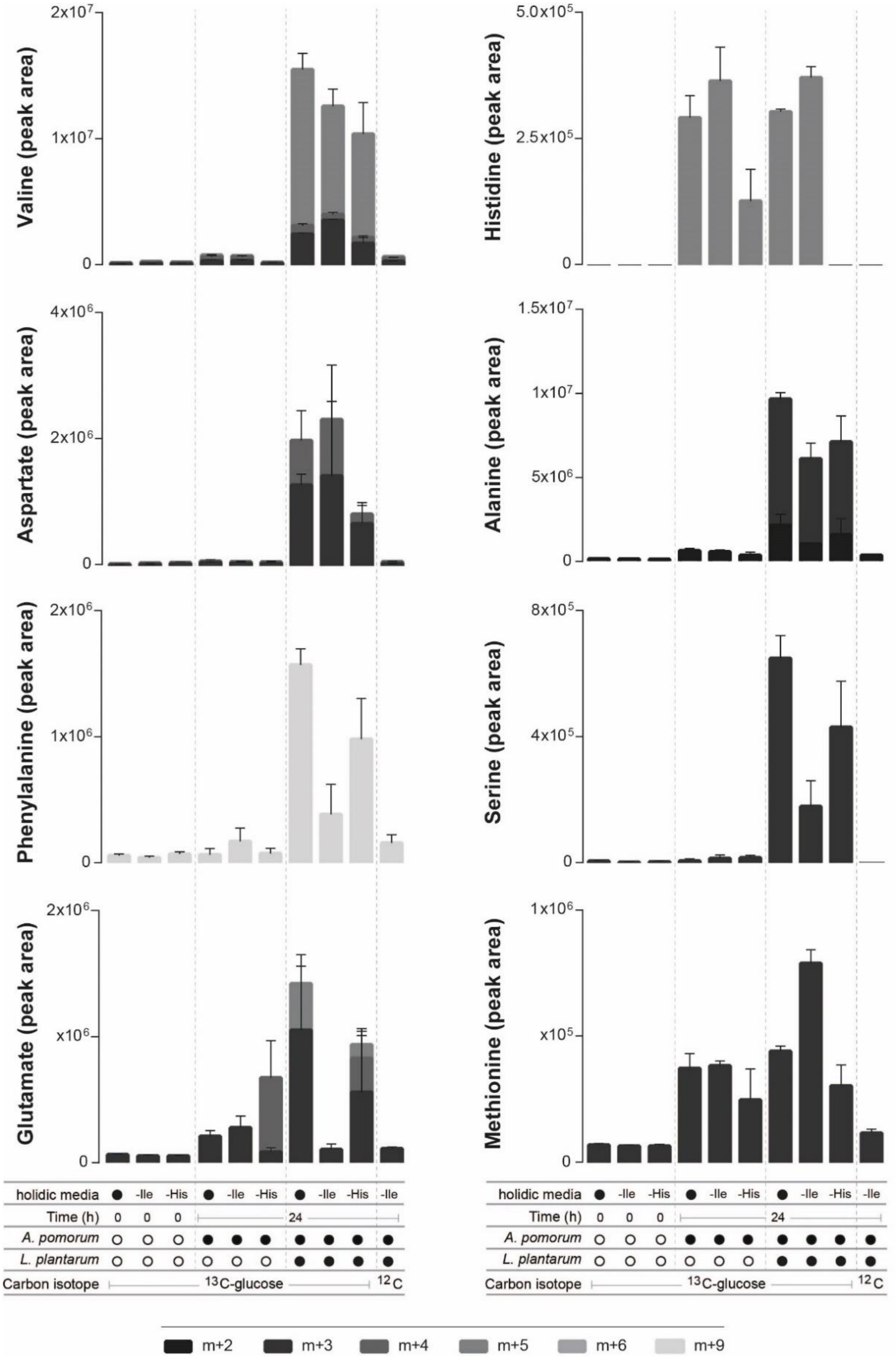


**Supplementary Figure 4 – Synthesis of AAs is increased in the presence of *Lp* (after 24h of growth).** The stacked bars represent the relative amount of isotopomers of each indicated AA measured in the supernatant of bacterial cultures grown in complete liquid holidic medium or in media lacking Ile or His containing uniformly $^{13}C$ labelled glucose ($^{13}C$-glucose). Co-cultures were also grown in media lacking Ile and containing, unlabelled glucose ($^{12}C$-glucose) as a control for unspecific labelling. Heavy labelled Ile isotopomers were measured using LC-MS in samples collected after 24h of bacterial growth and displayed as metabolite peak area. No bacteria were added in media representing time "0". The number of heavy carbons incorporated per Ile molecule is indicated as m+n, where n = the number of $^{13}C$. Filled black circles represent the presence of specific bacteria in the culture. Open black circles represent the absence of specific bacteria. Each data point represents the mean of three biological replicates and the error bars represent the standard error of the mean for each isotopomer.

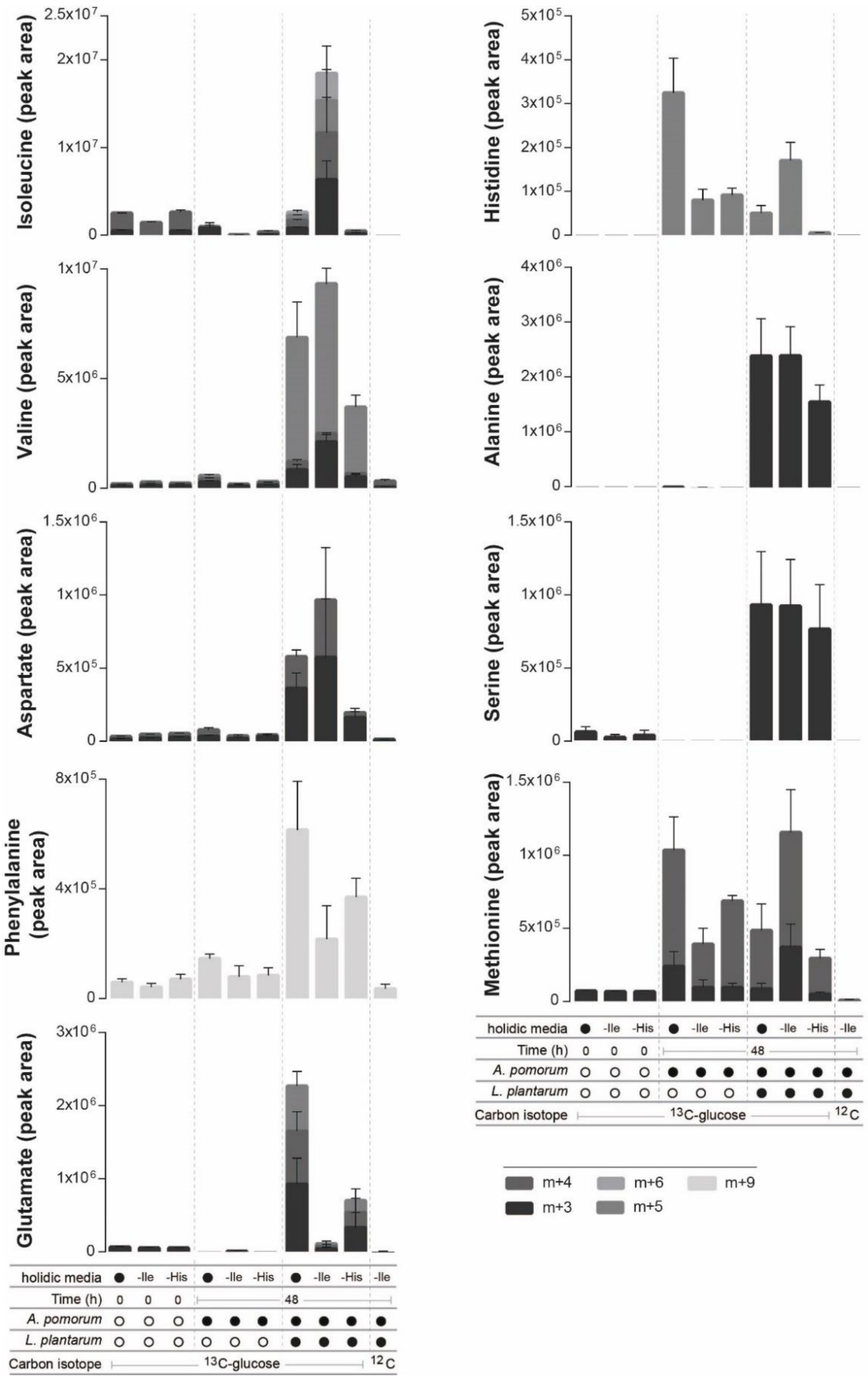


**Supplementary Figure 5 – Synthesis of AAs is increased in the presence of *Lp* (after 48h of growth).** The stacked bars represent the relative amount of isotopomers of each indicated AA measured in the supernatant of bacterial cultures grown in complete liquid holidic medium or in media lacking Ile or His containing uniformly $^{13}$C labelled glucose ($^{13}$C-glucose). Co-cultures were also grown in media lacking Ile and containing, unlabelled glucose ($^{12}$C-glucose) as a control for unspecific labelling. Heavy labelled Ile isotopomers were measured using LC-MS in samples collected after 48h of bacterial growth and displayed as metabolite peak area. No bacteria were added in media representing time "0". The number of heavy carbons incorporated per Ile molecule is indicated as m+n, where n = the number of $^{13}$C. Filled black circles represent the presence of specific bacteria in the culture. Open black circles represent the absence of specific bacteria. Each data point represents the mean of three biological replicates and the error bars represent the standard error of the mean for each isotopomer.

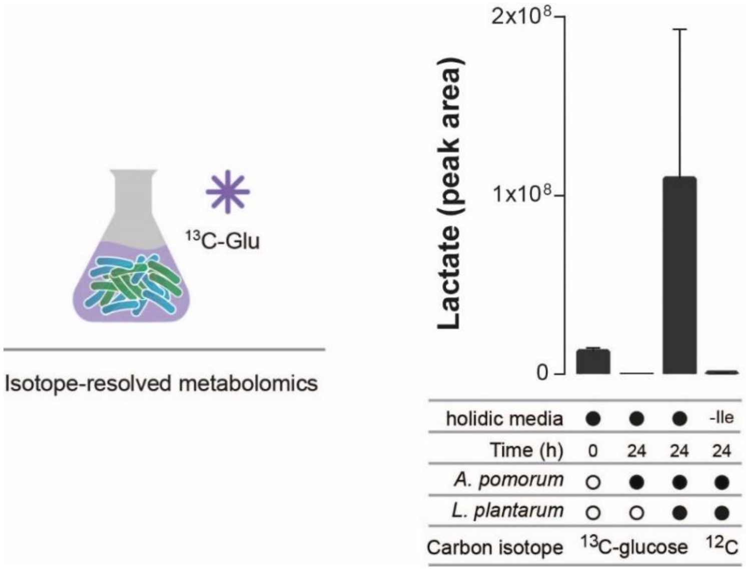


**Supplementary Figure 6 – Lactate is produced in *Lp* cultures.** Bars represent the relative amount of heavy lactate measured in the supernatant of bacterial cultures grown in complete liquid holidic medium containing uniformly $^{13}$C labelled glucose ($^{13}$C-glucose). Co-cultures were also grown in media lacking Ile and containing generic, unlabelled glucose ($^{12}$C-glucose) as a control for unspecific labelling. Heavy labelled lactate was measured using LC-MS in samples collected after 24h of bacterial growth and displayed as metabolite peak area. No bacteria were added in media representing time “0”. Only the m+3 isotopomer was detected. Filled black circles represent the presence of specific bacteria in the culture. Open black circles represent the absence of specific bacteria. Each data point represents the mean of three biological replicates and the error bars represent the standard error of the mean for each isotopomer.

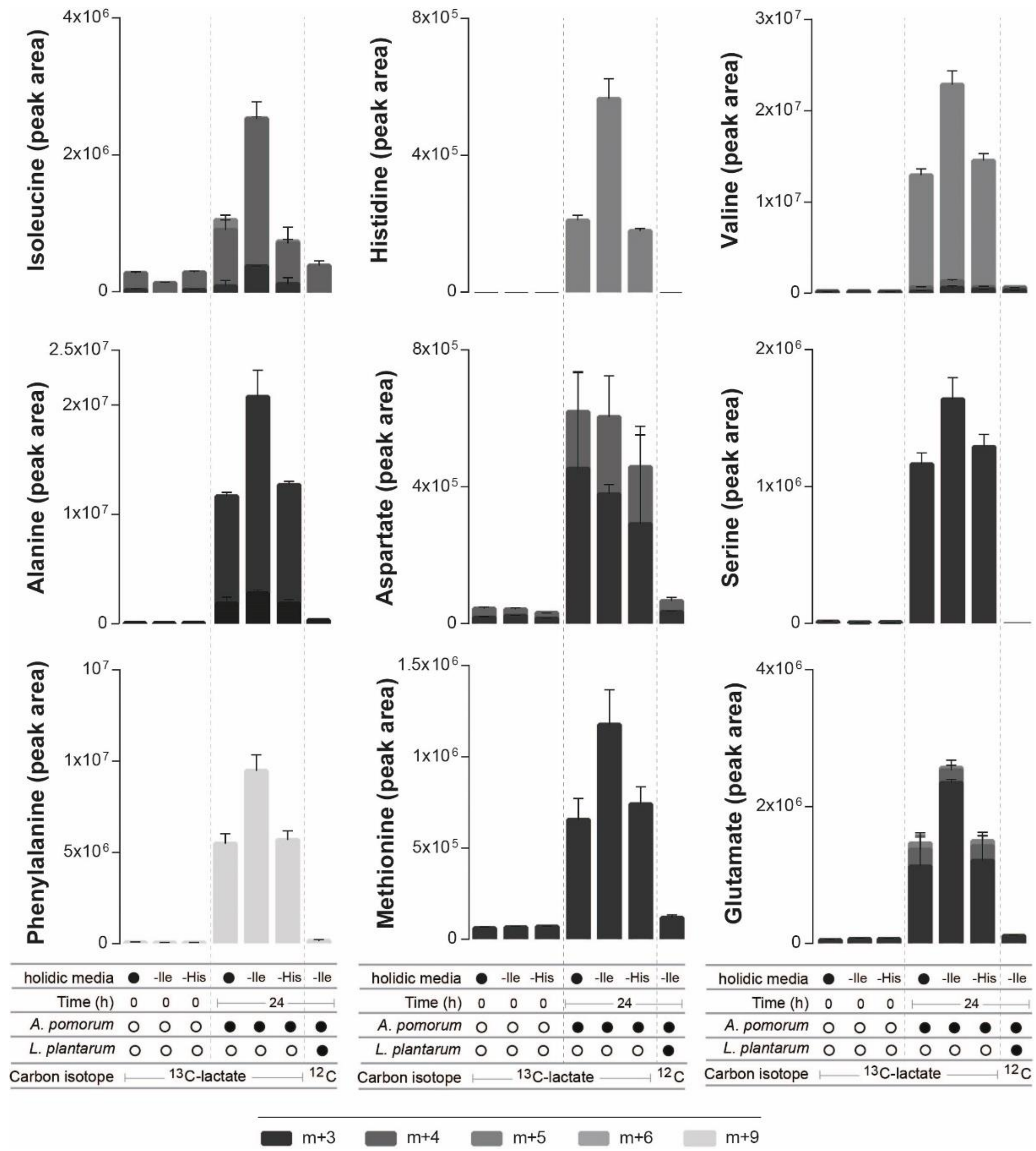


**Supplementary Figure 7 – Secreted AA synthetized from lactate (after 24h of growth).** The stacked bars represent the relative amount of isotopomers of each indicated AA measured in the supernatant of bacterial cultures grown in complete liquid holidic medium or in media lacking Ile or His containing uniformly $^{13}C$ labelled lactate ($^{13}C$-lactate, 20 g/l). Co-cultures were also grown in media lacking Ile containing unlabelled glucose ($^{12}C$-glucose) as a control for unspecific labelling. Heavy labelled Ile isotopomers were measured using LC-MS in samples collected after 24h of bacterial growth and displayed as metabolite peak area. No bacteria were added in media representing time "0". The number of heavy carbons incorporated per Ile molecule is indicated as m+n, where n = the number of $^{13}C$. Filled black circles represent the presence of specific bacteria in the culture. Open black circles represent the absence of specific bacteria. Each data point represents the mean of three biological replicates and the error bars represent the standard error of the mean for each isotopomer.

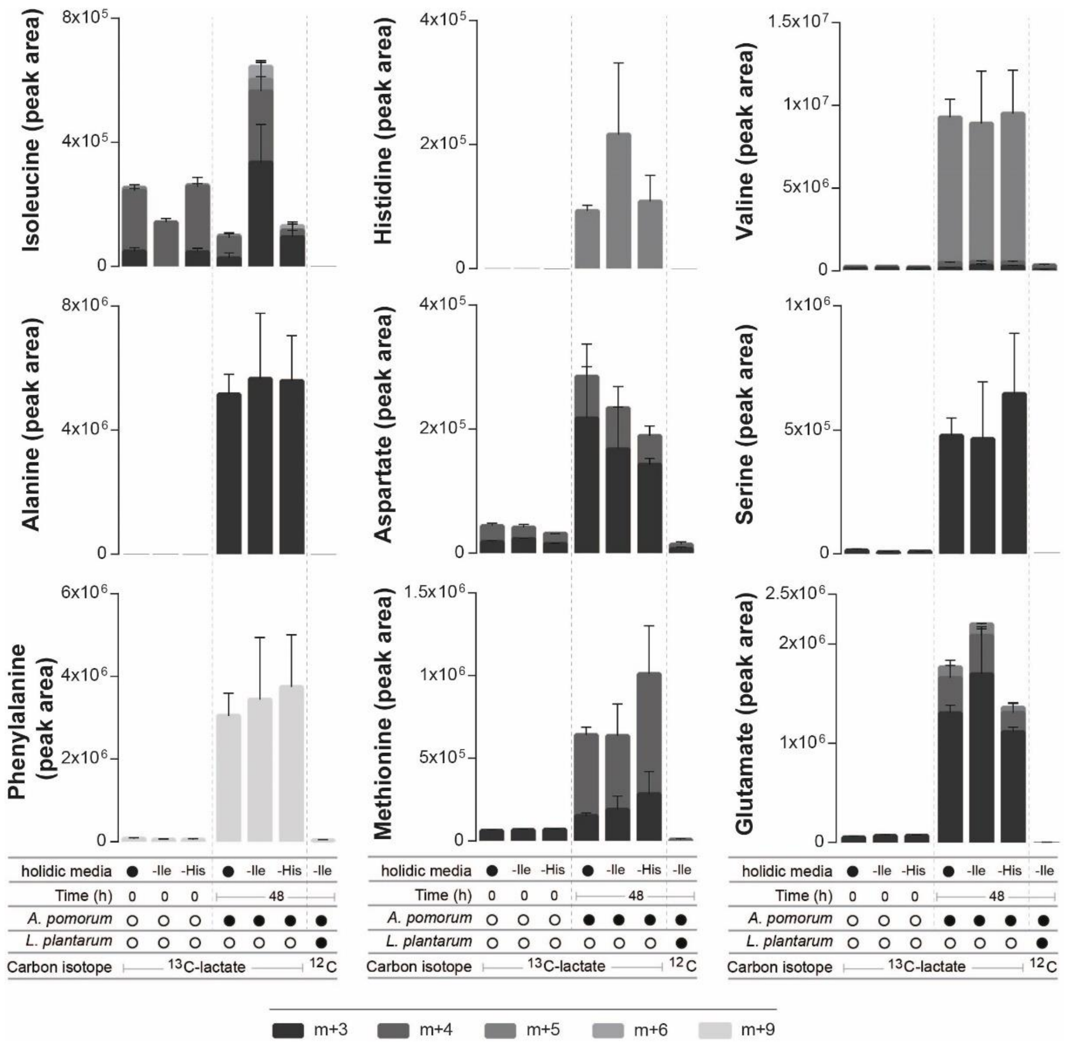


**Supplementary Figure 8 – Secreted AA synthetized from lactate (after 48h of growth).** The stacked bars represent the relative amount of isotopomers of each indicated AA measured in the supernatant of bacterial cultures grown in complete liquid holidic medium or in media lacking Ile or His containing uniformly $^{13}$C labelled lactate ($^{13}$C-lactate, 20 g/l). Co-cultures were also grown in media lacking Ile containing unlabelled glucose ($^{12}$C-glucose) as a control for unspecific labelling. Heavy labelled Ile isotopomers were measured using LC-MS in samples collected after 48h of bacterial growth and displayed as metabolite peak area. No bacteria were added in media representing time "0". The number of heavy carbons incorporated per Ile molecule is indicated as m+n, where n = the number of $^{13}$C. Filled black circles represent the presence of specific bacteria in the culture. Open black circles represent the absence of specific bacteria. Each data point represents the mean of three biological replicates and the error bars represent the standard error of the mean for each isotopomer.

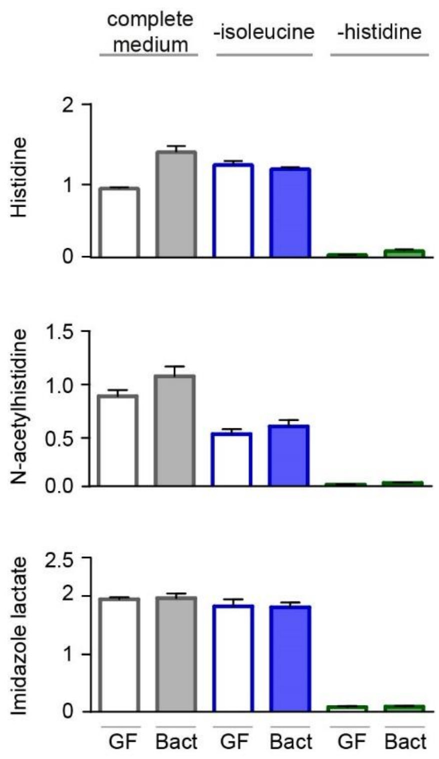


**Supplementary Figure 9 – His and His degradation products are not increased in heads of flies inoculated with bacteria.** Bars represent the relative abundance of His and His degradation metabolites as measured in metabolomics experiments from heads of germ-free flies (GF) and flies inoculated with commensal bacteria (Bact) and maintained in either complete holidic medium (grey boxes), medium lacking Ile (blue boxes) or medium lacking His (green boxes). Data are plotted as the mean of five biological replicates and the error bars represent the standard error of the mean.

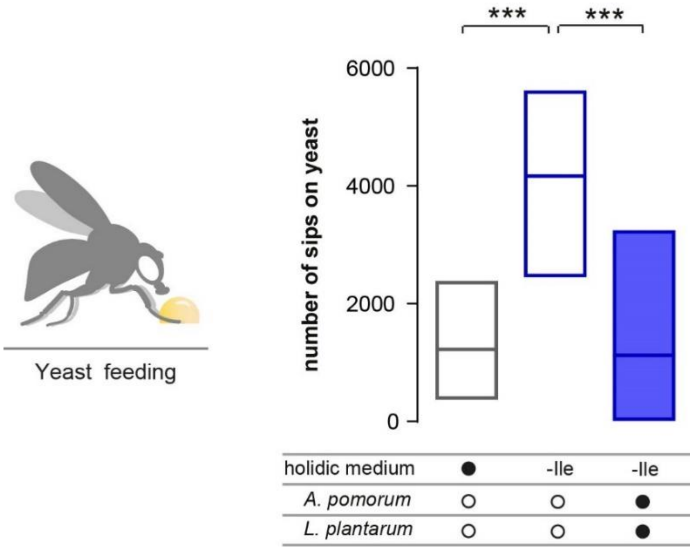


**Supplementary Figure 10 – Glucose and sucrose affect fly's yeast preferences to the same extent.** Number of sips on yeast of germ-free (empty) flies or flies biassociated (blue) with *Ap* and *Lp* that were kept in complete holidic medium or medium without Ile (-Ile) where an equimolar concentration of the monosaccharide glucose (100 mM) was used as a carbon source in the holidic medium recipe (see material and methods section) instead of the disaccharide sucrose (50 mM). Boxes represent median with upper and lower quartiles. n = 40-51. Significance was tested using a Kruskal–Wallis test with a Dunn's multiple comparison test. Filled black circles represent a complete holidic medium or association with *Ap*+*Lp*. Open black circles represent the absence of bacteria. *** $p \leq 0.001$.
